# Red Blood Cells Function as DNA Sensors

**DOI:** 10.1101/2021.02.11.430782

**Authors:** Metthew Lam, Sophia Murphy, Dimitra Kokkinaki, Alessandro Venosa, Scott Sherrill-Mix, Carla Casu, Stefano Rivella, Aaron Weiner, Jeongho Park, Sunny Shin, Andrew Vaughan, Beatrice H. Hahn, Audrey R. Odom John, Nuala J. Meyer, Christopher A. Hunter, G. Scott Worthen, Nilam S. Mangalmurti

## Abstract

Erythrocytes have long been mistaken as exclusively inert oxygen carriers lacking immune function. Here we show that red blood cells (RBCs) serve as immune sensors through surface expression of the nucleic acid-sensing toll-like receptor 9 (TLR9), a classically endosomal receptor that initiates immune responses following the detection of unmethylated CpG motifs present in pathogen and mitochondrial DNA. Mammalian RBCs express TLR9 on their surface and bind CpG-containing bacterial, malarial, and mitochondrial DNA. Erythrocyte-bound CpG DNA increases during infection, and CpG-carrying RBCs trigger accelerated erythrophagocytosis and innate immune activation characterized by RBC-TLR9 dependent local and systemic cytokine production. Thus, RBC nucleic acid detection and capture regulates red cell clearance and immune responses and provides evidence for RBCs as innate immune sentinels during pathologic states.

**One Sentence Summary:** The ability of RBCs to detect and bind cell-free nucleic acids contributes to immunity during acute inflammatory states.

---

Red blood cells (RBCs) comprise the majority of circulating cells in mammals and are essential for respiration. Although non-gas exchanging functions of the red cell such as chemokine regulation, complement binding, and pathogen immobilization have been described, RBC immune function remains enigmatic ^1-3^. RBCs transit through all tissues and contact pathogen and self-derived inflammatory mediators in the circulation, positioning them as ideal messengers between distant organs. Indeed, we have recently demonstrated that RBCs bind and scavenge nucleic acids away from the lung during basal conditions at homeostasis ^4^, yet the role of RBC-nucleic acid-binding in the host immune response during inflammation is unknown.

Nucleic acid-sensing is an essential function of the innate immune system necessary to detect infection and sterile injury ^5^. Evolutionarily conserved nucleic acid-sensing toll-like receptors (TLRs) identify nucleic acids derived from self and pathogens and play a central role in inflammation by promoting inflammatory cytokine secretion, immune cell maturation, and proliferation ^5-10^. Elevated cell-free CpG-containing DNA is a hallmark of infection and sterile injury ^7,8,11^. Common to these inflammatory pathologies is acute anemia, a significant cause of morbidity observed during sepsis and critical illness. Several studies have suggested a role for intracellular nucleic acids and TLR signaling in monocyte/macrophages in developing inflammatory anemia and cytopenias, yet whether RBCs themselves and cell-free DNA contribute to red cell clearance is unknown ^12-14^.

We recently discovered that RBCs express intracellular TLR9 and scavenge cell-free (CpG-containing) mitochondrial DNA (cf-mtDNA) under homeostatic conditions ^4^. These data suggested that TLR9-mediated sequestration of cf-mtDNA by RBCs represents a protective mechanism that clears toxic cell-free nucleic acids from the circulation in healthy hosts ^4^. However, how RBC-dependent CpG-binding contributes to the inflammatory response during infection remains unknown. Here we show that TLR9 is expressed on the RBC surface and that DNA binding by RBC TLR9 during inflammatory states leads to accelerated RBC clearance and systemic inflammation, thus linking RBC nucleic acid binding with innate immune activation during inflammatory states.

## Results

### TLR9 is increased on the surface of RBCs during sepsis, and RBCs bind pathogen DNA

Although TLR9 is an endosomal nucleic acid-sensing receptor, recent studies have identified the presence of TLR9 on the surface of intestinal epithelial cells, splenic dendritic cells, platelets, and a minor fraction of peripheral blood mononuclear cells (PBMCs) ^15-18^. We have previously detected intracellular TLR9 in RBCs but were unable to detect surface expression of TLR9 using permeabilization-dependent antibodies ^4^. However, when we used antibodies to a larger epitope in the TLR9 ectodomain, TLR9 was readily detected on intact non-permeabilized human and murine RBCs (Fig. 1A). We verified RBC TLR9 expression using confocal microscopy (Fig. 1B). We also asked whether RBCs from non-human primates express surface TLR9. Chimpanzee RBCs, like their human counterparts, express surface TLR9 (fig. S1). Thus, TLR9 expression is conserved in mammalian RBCs with the DNA-binding ectodomain on the RBC surface.

**Figure 1.**
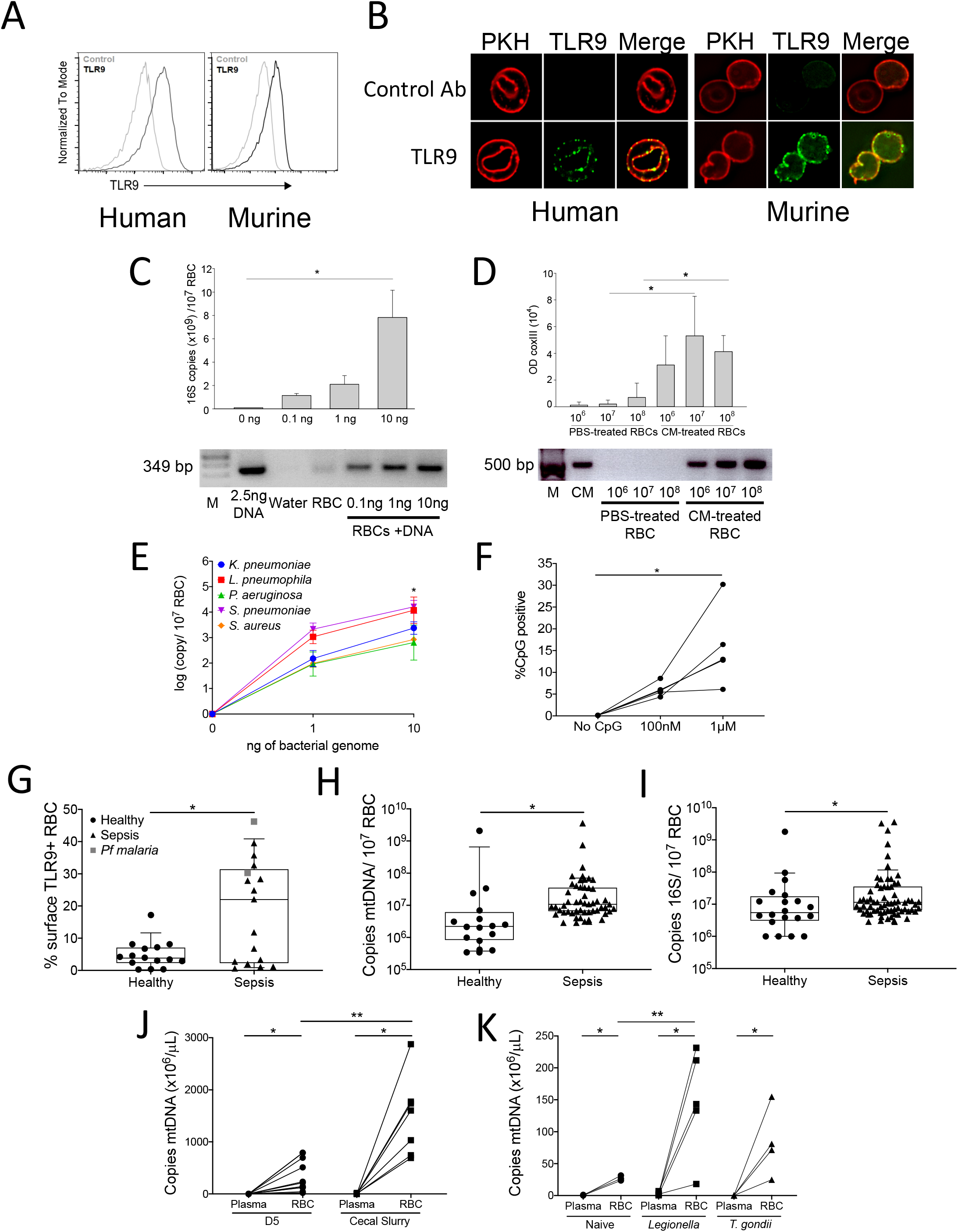
DNA binds mammalian erythrocytes through surface expressed TLR9. (**A**) Non-permeabilized, erythrocytes from a healthy human donor and mouse were labeled with TLR9 ectodomain specific antibodies and analyzed by flow cytometry. (**B**) Confocal images showing surface expression of TLR9 and PKH 26-labeled human and murine red cell membranes. (**C**) RBCs from healthy human donors were incubated with increasing doses of *Legionella* DNA (0.1, 1, 10 ng), and the presence of bacterial DNA was measured by quantitative PCR of 16S amplification (top panel). 3 individual studies, \**P*=0.008, one-way ANOVA. Amplicons from one representative study are shown (bottom panel). Lane 1:M=marker. (**D**) Increasing amounts of RBCs (10^6^, 10^7^, 10^8^) were incubated with PBS or *P. falciparum*-positive culture medium (CM), and parasite DNA binding was quantified by DNA extraction and *P. falciparum* mtDNA amplification (COX III). Optical density (OD) quantification of *P. falciparum* DNA bound to RBCs is shown (top panel). *P*=NS for 10^6^ RBCs, \**P*<0.04 for 10^7^ RBCs, and \**P*<0.02 for 10^8^ RBCs, t-test comparing PBS v CM treated RBCs performed. RBCs from 3 individual donors were tested in 3 independent experiments. Amplicons from one representative study (bottom panel) is shown. Lane 1: marker. (**E**) Human RBCs binding to increasing doses of bacterial genomic DNA, and bacteria-specific qPCR primers were used to quantify RBC-associated DNA. n=4 healthy donors. * *P*<0.05 compared to 0 ng, Kruskal-Wallis test, Dunn’s post-hoc analysis. (**F**) CpG-binding of RBCs from 5 healthy donors were tested with 2 doses of CpG. n=5 donors, \**P*=0.002, Kruskall-Wallis test, Tukey post-hoc analysis. (**G** – **I**) Comparison of healthy donor and septic patient RBC, n >15. (G) Surface TLR9 expression on RBC. *\*P*=0.041, Mann Whitney U-test. Patients with *P. falciparum*-induced sepsis are denoted by black circles. Quantification of (H) mitochondrial DNA and (I) 16s DNA on RBCs from healthy donors and patients with sepsis, as determined by quantitative PCR, are compared \**P*<0.001, n=16 healthy and 48 sepsis for mtDNA;*\*P*=0.023 for 16s, n=20 healthy and 64 sepsis for 16s. Each dot represents a different patient. Mann Whitney U-test for all comparisons. (**J**) Mitochondrial DNA (*mtCo1*) on plasma and RBCs from mice 6 hours following cecal slurry injection, \**P*<0.001, paired t-test for RBCs v plasma *mtCo1* in D5W group, *\*P*=0.002 paired t-test for RBCs v plasma for CS injected mice, *\*P*=0.003 for RBC mtDNA, D5W v CS injected mice, t test; n=7-11 mice, three independent studies. (**K**) *mtCo1* on RBC and plasma of *L. pneumophilia*- or *T. gondii*-infected C57/B6 mice. *\*P*<0.008 for *L. pneumophilia* plasma v RBCs, \**P*=0.029 for *T. gondii* plasma v RBCs, paired t-test. \*\**P*=0.053 for naïve RBCs v *L. pneumophilia* RBCs.

Immunostimulatory unmethylated CpG motifs are a feature of microbial DNA; we, therefore, asked whether RBCs could detect pathogen DNA. The ability of RBCs from healthy human donors to bind bacterial DNA or malarial mtDNA was tested by incubating RBCs with genomic DNA from *Legionella pneumophilia* or media from *Plasmodium falciparum* erythrocyte culture. Following incubation with bacterial DNA or *P. falciparum* DNA, RBCs were isolated, and PCR for the 16s ribosomal RNA gene (bacterial DNA) or coxIII (malarial mtDNA) was performed. We found a dose-dependent increase in amplifiable microbial DNA on RBCs following incubation with bacterial or malarial DNA (Fig. 1C and D). We confirmed human RBCs’ ability to bind bacterial and malarial DNA using genomic DNA from common pathogens and synthetic CpG based on sequences found in the *P. falciparum* genome (Fig. 1E and F) ^19^. Therefore, human RBCs can bind pathogen DNA.

The TLR9 ligand, CpG-containing mitochondrial DNA, is elevated in the circulation during sepsis, a deadly syndrome defined by the dysregulated host response to infection ^11,20,21^. We therefore examined TLR9 expression on RBCs from critically ill patients with sepsis. RBCs were prospectively collected from patients enrolled in a cohort designed to study sepsis (Molecular Epidemiology of SepsiS in the ICU, MESSI cohort) at the University of Pennsylvania. Flow cytometry for TLR9 was performed on intact non-permeabilized RBCs. We found that surface TLR9 was increased on RBCs from patients with sepsis compared with RBCs from healthy donors (Fig. 1G). We next performed quantitative PCR (qPCR) to determine the mtDNA content of RBCs obtained from healthy volunteers and critically ill patients with sepsis ^4,11^. We found that mtDNA was increased on RBCs from patients with sepsis compared with RBCs from healthy controls (Fig. 1H). Because microbial DNA is a potent activator of TLR9 and our data demonstrate RBC detection of microbial DNA, we measured 16s content on RBCs by qPCR. Consistent with our *in vitro* findings of bacterial DNA acquisition by RBCs, 16s is elevated on RBCs during human sepsis (Fig. 1I). Thus, TLR9 and its ligand, CpG-containing DNA, are elevated on the surface of RBCs during human sepsis.

We next asked whether CpG-containing host mitochondrial DNA bound RBCs during infection. We measured mitochondrial DNA (coxI) in the plasma and on RBCs in a murine model of sepsis and murine models of bacterial pneumonia and systemic parasite infection (*L. pneumophila and Toxoplasma gondii*). As seen in Figure 1J and K, mtDNA was elevated on RBCs when compared with plasma in all animal models of infection. These data demonstrate that CpG-containing mtDNA is sequestered on RBCs during pneumonia, parasitic infection, and polymicrobial infection ^4^.

### DNA binding results in altered RBC structure and function

During sterile inflammation and infection, cell-free CpG-containing mtDNA is elevated in the plasma ^9,11^. Because altered RBC morphology is a common feature of sepsis and critical illness, we examined the effect of excess cell-free CpG on RBC morphology and function ^22-26^. Extensive alterations of RBC morphology following high dose CpG treatment (100 nM) were visualized by electron microscopy (Fig. 2A and B). RBC shape changes were associated with marked differences in the distribution of the cytoskeletal proteins spectrin and actin following CpG binding (Fig. 2C). We next examined the distribution of the RBC membrane protein Band 3 following CpG DNA treatment. As a control, we used GpC DNA, which binds TLR9 without causing TLR9 activation or conformational changes ^27^. As shown in Figures 2D and E, treatment with CpG DNA, but not GpC DNA, led to alterations in Band 3 distribution. Taken together, these data indicate that CpG DNA binding alters RBC morphology in a TLR9-dependent manner. We next examined the morphology of RBCs following CpG incubation using imaging flow cytometry. This analysis identified a population of RBCs with aberrant morphology following incubation with CpG. The automated feature finder was utilized to discriminate RBC subpopulations based on morphology, and mean pixel and intensity features were identified to best differentiate smooth from altered cells (Fig. 2F), the altered cells were all TLR9 positive (Fig. 2G). In the presence of low levels of extracellular CpG DNA, RBCs remained morphologically unaltered.

**Figure 2.**
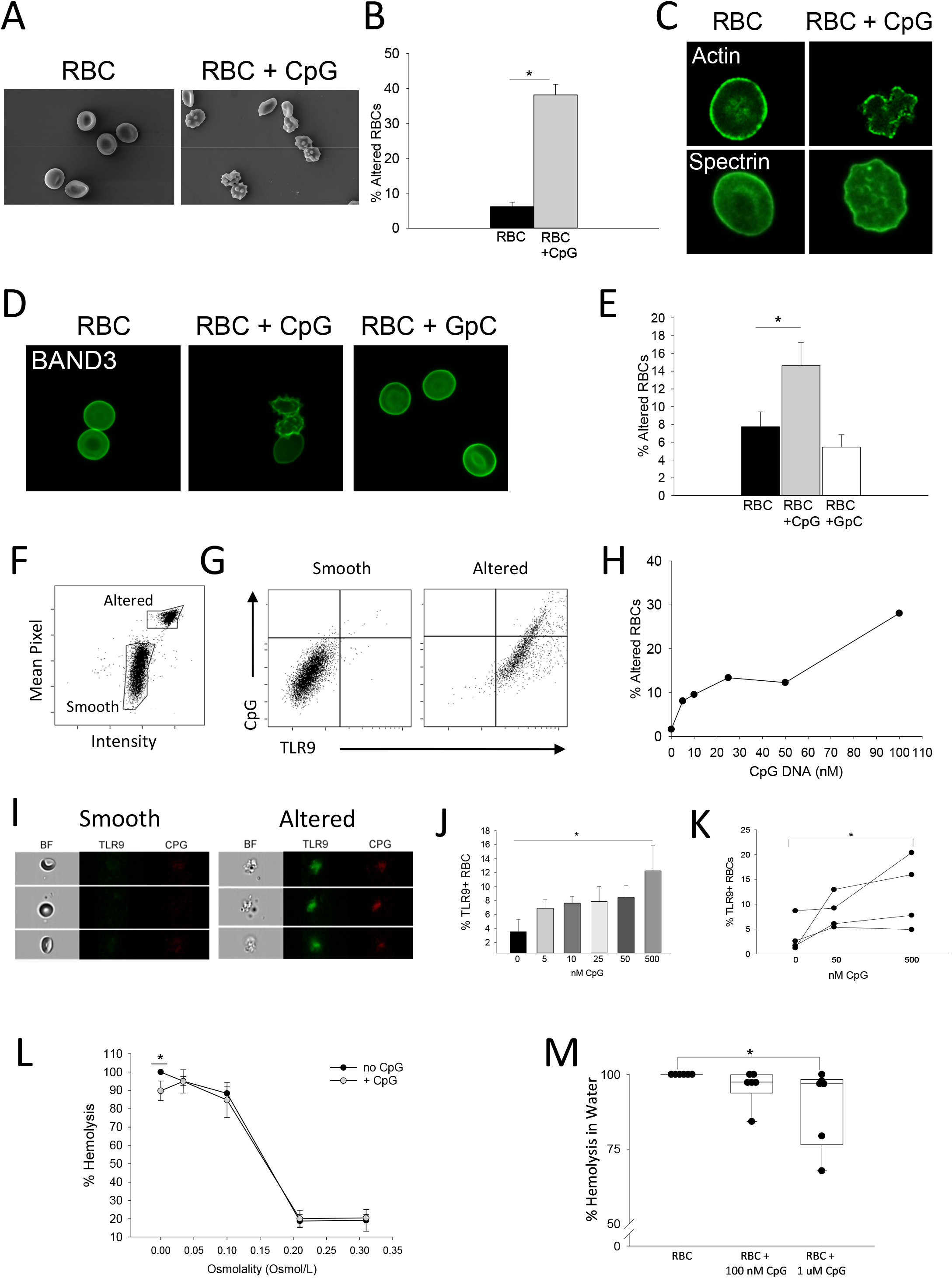
Human RBCs undergo structural alterations upon CpG binding. (**A**) Scanning electron microscopy of human RBCs following CpG incubation. (**B**) Quantification of RBC alteration in untreated and CpG-treated RBCs observed by electron microscopy. Alteration is defined by loss of biconcave disk shape and formation of echinocytes. Cells from 5 separate fields were counted and averaged. \**P*<0.001, t-test. (**C**) Confocal images showing RBC cytoskeletal proteins actin and spectrin following CpG treatment. (**D**) Confocal imaging displays Band 3 distribution following CpG and control GpC treatments. (**E**) Quantification of RBC alteration by Band 3 staining displays CpG treated cells are more altered than GpC treated cells and untreated cells. RBCs from 4 individual donors tested. \**P*=0.033, one-way ANOVA with Dunn’s post-hoc analysis. **(F-I)** Imaging flow cytometry analysis on CpG-treated human RBCs. (F) Imaging flow cytometry reveals smooth and altered RBC populations as defined by Mean Pixel and Intensity parameters. (G) Smooth and altered RBC populations were analyzed for CpG binding and TLR9 expression. (H) Percent of altered RBC in increasing doses of CpG. (I) Images of smooth and altered RBCs displaying differences in CpG and surface TLR9 detection. (**J-K**). Surface TLR9 detection on RBC treated with increasing doses of CpG, n=4. (J) Summary statistics and (K) results from individual donors \**P*=0.052 (0 nM v 500 nM CpG, paired t-test). (**L**) Osmotic fragility of healthy human RBCs pre-treated with PBS, 100 nM CpG, or 1 uM CpG. (**M**) Hemolysis of RBCs pre-treated with PBS, 100 nM CpG, or 1 uM CpG after incubating cells in water. RBCs from 6 independent donors were tested. \**P*=0.02, Kruskal-Wallis test, Dunn’s post-hoc analysis \**P*=0.022 no CpG v 1uM, *P*=0.132 no CpG v 100 nM.

However, addition of increasing amounts of CpG resulted in malformed RBCs (Fig. 2H), representative images of smooth and altered cells are shown in Figure 2I. Because we observed increased surface TLR9 accessibility in patients with sepsis, we examined RBC surface TLR9 expression following treatment with diverse stimuli. LPS, TNF and PMA did not alter surface TLR9 detection whereas CpG and lipid disrupting agents (hydrogen peroxide) increased surface TLR9 positivity (Fig. 2J and K and fig, S2).

We next examined the effect of excess cell-free CpG on RBC function by measuring osmotic fragility of human RBCs and viability. As shown in figures 2L and M, the addition of CpG resulted in reduced osmotic fragility. Collectively, these data suggest that surface accessibility of the TLR9 ectodomain is tunable by CpG exposure and that CpG binding to RBCs alters RBC structure and function.

### CpG binding by RBCs leads loss of CD47 detection

RBC survival is determined by multiple factors, including membrane integrity, phosphatidylserine (PS) externalization, and CD47 expression. RBC viability was assessed by loss of membrane integrity using the dye calcein-AM. As seen in supplemental figure 3A and B, CpG did not lead to a loss of RBC membrane integrity. Because PS externalization serves as an “eat me” signal and CD47 expression serves as a self-preservation or “don’t eat me” signal, we examined both PS externalization and CD47 expression following CpG treatment of RBCs. While we did not observe an increase in PS positive cells, we observed loss of CD47 as measured by binding of the antibody CC2C6, which detects the antiphagocytic epitope of CD47 (fig. S3C and D and Fig. 3A and B). We observed that cells that were CC2C6 negative following the addition of CpG DNA had bound higher amounts of DNA than those that were CC2C6 positive (fig. S4A), suggesting that CpG-acquisition by RBCs led to loss of CD47 detection. These findings were confirmed with imaging flow cytometry which revealed the majority of CD47 dim cells were altered (fig. S4).

**Figure 3.**
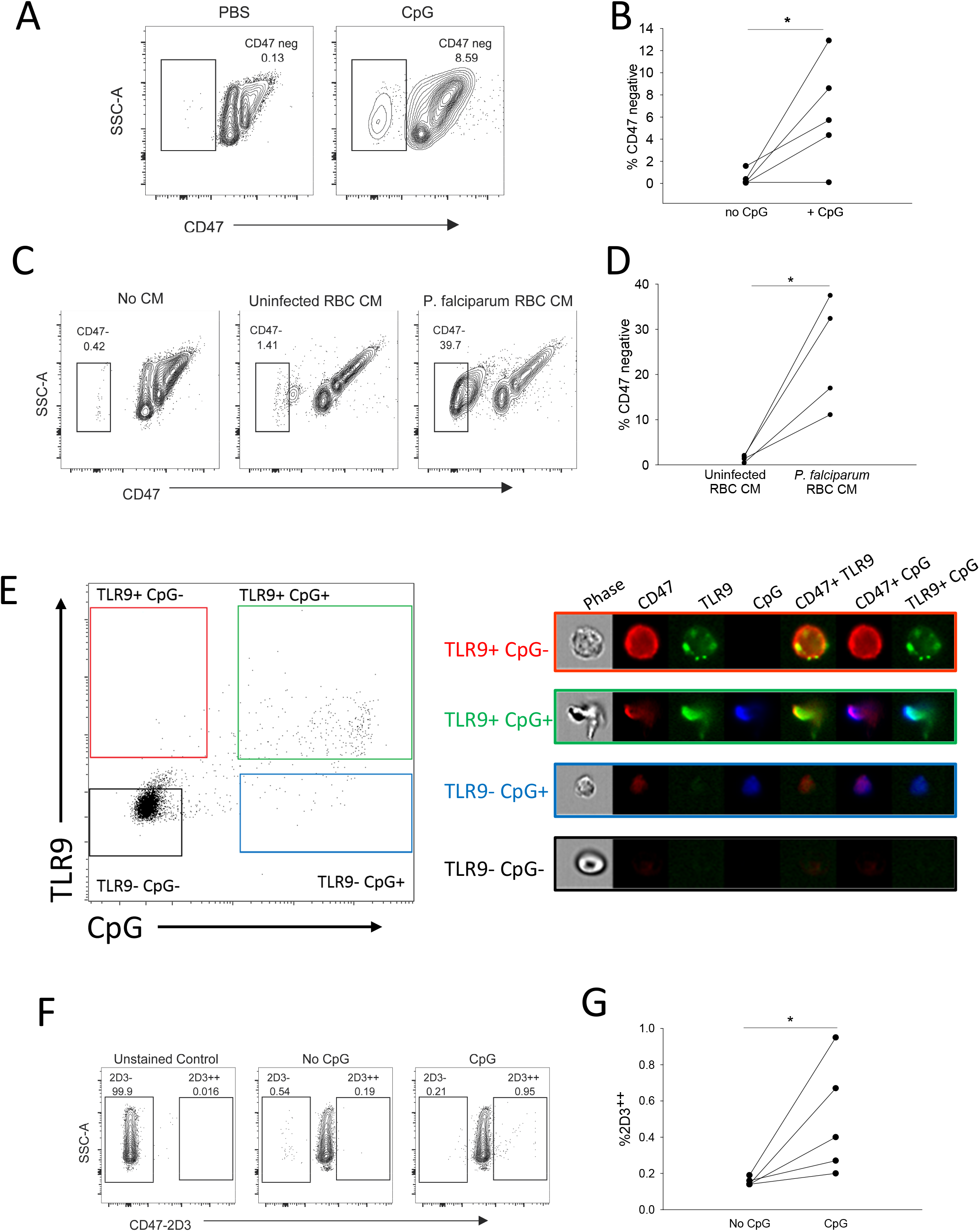
DNA binding by surface expressed TLR9 induces RBC loss of self. (**A**) Representative results for CD47 detection (with antibody clone CC2C6) on human RBC following incubation with 10μg/mL CpG and (**B**) the corresponding results from five donors. \**P*=0.04, paired t-test (**C**) CD47 detection in human RBC treated with condition medium derived from *P. falciparum*-infected RBC culture and (**D**) the corresponding results from 4 donors. *\*P*=0.031, paired t-test. (**E**) Imaging flow cytometry on CpG-treated human RBC probed for CD47 and TLR9. (**F**) Induction of conformational change in CD47 by CpG as indicated by detection of anti-CD47 (clone 2D3), a damaged-associated conformational epitope. One representative flow analysis is shown and (**G**) the corresponding quantification data form 5 individual donors \**P*=0.038 by t-test, P=0.06 by paired t-test.

Because naïve RBCs bind malarial DNA, we asked if malarial DNA would induce loss of the CD47 antiphagocytic epitope in uninfected erythrocytes from healthy human donors. CpG sequences, based on sequences found in the *P. falciparum* genome, led to a loss of CD47 detection on RBCs obtained from healthy donors (fig. S5C and D) ^19^. Incubation of naïve human RBCs with *P. falciparum* culture supernatants led to robust loss of the antiphagocytic epitope (Fig. 3C and D).

CD47 associates with the Band 3 complex, a macro-complex of proteins in the RBC membrane ^28,29^. We therefore asked whether CD47 was also in complex with TLR9. We found that these two proteins co-immunoprecipitated (Fig. S5A). We confirmed this physical interaction using confocal microscopy, which revealed co-localization of TLR9 and CD47 in the RBC membrane (Fig. S5B). Because we had observed morphological alterations in the RBC membrane following CpG binding, we performed imaging flow cytometry to better characterize the distribution of surface TLR9 and CD47 in intact RBCs following CpG DNA addition. We incubated RBCs with 100nM CpG and found that CpG DNA led to alterations in RBC structure as well as CD47 and TLR9 redistribution (Fig. 3E). TLR9-positive, CpG-negative cells demonstrated punctate surface TLR9 staining and uniform surface CD47 distribution, whereas TLR9-positive, CpG-positive cells showed alteration of the membrane and clustering of TLR9 and CD47. These findings suggest that a conformational change of CD47 occurs following CpG-binding by RBCs.

We next asked whether conformational changes of CD47 were associated with the altered localization observed after CpG binding. Conformational changes in CD47 can be detected by an increase in binding of the anti-CD47 antibody 2D3 ^30^. This antibody detects an epitope on CD47 that has undergone conformational changes and is present on “damaged”, experimentally aged, and sickle RBCs ^30,31^. Incubation of RBCs with CpG for 2 hours led to increased detection of this CD47 epitope using the 2D3 antibody (Fig.3F and G).

### DNA-carrying RBCs undergo accelerated erythrophagocytosis and initiate innate immune responses

Because CD47 is a marker of self, and loss of CD47 leads to accelerated erythrophagocytosis by red pulp F4/80 positive splenic macrophages (RPM) ^32^, we asked whether CpG binding by RBCs would result in accelerated clearance of RBCs by RPM. GFP-expressing RBCs were treated with PBS or CpG DNA for 2 hours before being washed and infused into mice. Analysis of spleens one hour following infusion revealed that F4/80-high macrophages ingested GFP RBCs (gating in fig. S6). Mice that received CpG-treated RBCs demonstrated a significantly higher percentage of erythrophagocytic macrophages than mice that received PBS-treated RBCs (Fig. 4A-C). Spleen weights of animals receiving CpG-treated RBCs were also elevated at 20 hours post-infusion compared with animals that received PBS-treated RBCs, consistent with increased erythrophagocytosis and splenic congestion (Fig. 4D). Collectively, these findings demonstrate that exposure of RBCs to high concentrations of CpG DNA leads to accelerated erythrophagocytosis *in vivo*.

**Figure 4.**
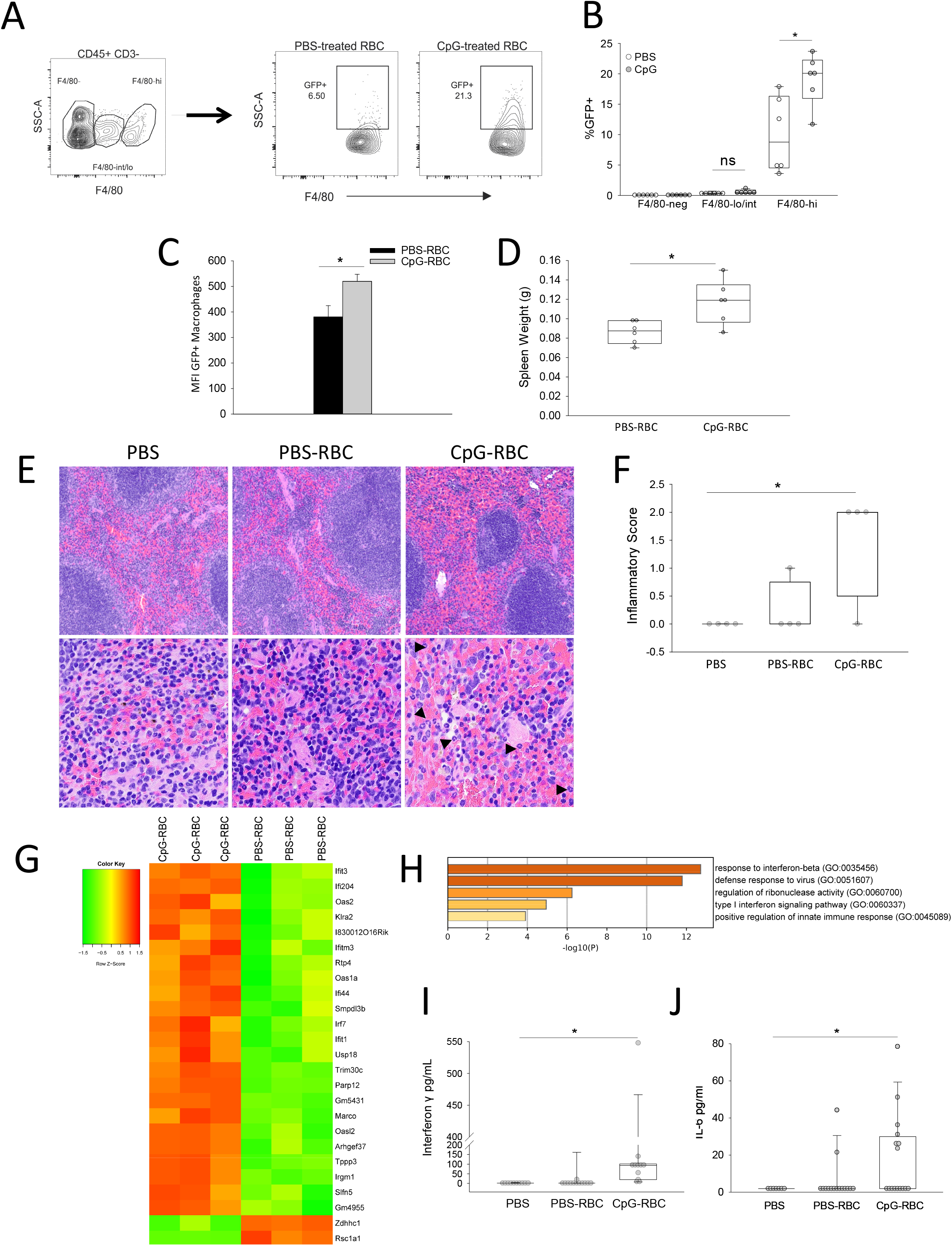
CpG-carrying RBCs undergo accelerated erythrophagocytosis and induce systemic inflammatory responses in naïve mice. (**A-D**) GFP-expressing RBC obtained from GFP-expressing mice were treated with PBS or CpG for 2hr at 37C and transfused to WT mice. 1hr post-infusion, CD45+ CD3-splenocytes were analyzed. Erythrophagocytosis by F4/80+ cells was investigated, n=6/group, 2-independent experiments. (A) Gating strategy for F4/80-positivity and subsequent GFP-positivity in target cells. (B) Erythrophagocytosis by cells expressing different levels of F4/80, splenic red pulp macrophages (RPM) are defined as CD45+ CD3-F4/80-hi, two-way ANOVA, Sidak’s multiple comparisons test. *P<0.0001 (C) MFI of GFP in RPM. \**P*=0.023, t-test. (D) Spleen weight of infused mice at 20hr post-infusion \**P*=0.013, t-test. (**E**-**J**) Physiological changes at 6 h in mice infused with PBS, PBS-treated RBC, or CpG-treated RBC. (E) H&E stained spleens sections revealing red pulp congestion and the presence of neutrophils, arrowheads indicate neutrophils in the CpG-RBC treated group. (F) Quantification of spleen injury, 3-4 mice/group from 2 independent studies is shown, \**P*=0.02, ANOVA with post-hoc Holm-Sidak, multiple comparisons. (G) Heatmap for RNA-seq analysis of spleens from WT mice infused with PBS- or CpG-treated RBCs after 6hr and (H) GO-term analysis of the top 25 differentially expressed genes. Quantification of plasma (I) IFNγ \**P*<0.001, n=10-12 mice/group and (J) IL-6 \**P*=0.036, n=7-16 mice/group, Kruskal-Wallis test.

Given our findings of accelerated clearance of DNA-treated RBCs in mice, we asked whether mtDNA and microbial DNA content on RBCs would differ between anemic septic patients and non-anemic septic patients. A hemoglobin threshold of 7 g/dL was used to define anemia based on the current standard of care and transfusion guidelines for septic critically ill patients ^33,34^. As seen in Table 1, anemic septic patients had higher RBC-associated mtDNA, 16S, and total (mtDNA + 16S) DNA than non-anemic septic patients supporting the *in vivo* and *in vitro* findings of accelerated erythrophagocytosis of DNA carrying RBCs.

**Table 1.** Differentially expressed genes.

| Top 25 DEG based<br>on EdgeR | unadjusted p-value from<br>EdgeR | adjusted p-<br>values from<br>EdgeR | unadjusted p-value from<br>DEseq2 | adjusted p-<br>values from<br>DEseq2 |
| --- | --- | --- | --- | --- |
| <u>Upregulated genes</u> |  |  |  |  |
| Ifit3 | 3.53E-04 | 5.11E-01 | 7.84E-07 | 6.86E-04 |
| Ifi204 | 1.20E-04 | 5.08E-01 | 7.54E-10 | 3.58E-06 |
| Oas2 | 5.26E-04 | 5.36E-01 | 1.49E-09 | 5.30E-06 |
| Klra2 | 1.02E-03 | 5.89E-01 | 1.17E-05 | 5.94E-03 |
| Ifit3b | 6.97E-04 | 5.36E-01 | 1.46E-06 | 1.09E-03 |
| Ifitm3 | 9.84E-04 | 5.89E-01 | 1.68E-05 | 7.72E-03 |
| Rtp4 | 6.70E-04 | 5.36E-01 | 4.85E-07 | 4.61E-04 |
| Oas1a | 7.68E-04 | 5.36E-01 | 7.36E-06 | 4.77E-03 |
| Ifi44 | 7.94E-04 | 5.36E-01 | 3.09E-07 | 3.38E-04 |
| Smpdl3b | 8.51E-04 | 5.36E-01 | 4.43E-06 | 3.00E-03 |
| Irf7 | 4.85E-04 | 5.36E-01 | 1.66E-08 | 3.93E-05 |
| Ifit1 | 2.04E-04 | 5.08E-01 | 3.27E-08 | 6.65E-05 |
| Usp18 | 6.09E-04 | 5.36E-01 | 9.32E-07 | 7.37E-04 |
| Trim30c | 7.24E-04 | 5.36E-01 | 1.45E-05 | 6.87E-03 |
| Parp12 | 2.46E-04 | 5.08E-01 | 3.48E-07 | 3.54E-04 |
| Gm5431 | 4.67E-04 | 5.36E-01 | 2.48E-07 | 2.99E-04 |
| Marco | 1.68E-04 | 5.08E-01 | 4.05E-08 | 7.21E-05 |
| Oasl2 | 3.47E-05 | 4.56E-01 | 3.32E-15 | 4.73E-11 |
| Arhgef37 | 6.29E-05 | 4.56E-01 | 4.25E-10 | 3.03E-06 |
| Tppp3 | 2.21E-04 | 5.08E-01 | 4.18E-09 | 1.19E-05 |
| Irgm1 | 8.33E-04 | 5.36E-01 | 1.21E-05 | 5.94E-03 |
| Slfn5 | 3.50E-04 | 5.11E-01 | 1.43E-07 | 2.26E-04 |
| Ifi206 | 6.99E-04 | 5.36E-01 | 9.79E-06 | 5.16E-03 |
| <u>Downregulated genes</u> |  |  |  |  |
| Zdhhc1 | 3.04E-04 | 5.11E-01 | 8.59E-01 | 1.00 |
| rsc1a1 | 5.39E-04 | 5.36E-01 | not found | not found |

We next examined whether CpG-carrying RBCs would alter innate immune responses. To isolate RBCs’ role in initiating systemic inflammation, we utilized a reductionist model of CpG-carrying RBC infusion. Splenic histology 6 hours following CpG-RBC (but not PBS-treated RBCs) infusion also revealed increased neutrophil infiltration and enhanced red pulp congestion (Fig. 4E and F). To further characterize the immune response following CpG-RBCs, we performed RNA-seq on spleens from mice treated with RBCs or CpG-treated RBCs. Table 2 lists the top 25 differentially expressed genes. CpG-RBCs elicited a transcriptomic response characterized by increased expression of interferon signaling pathway genes compared with PBS-treated RBCs (Fig. 4G and H). Given the central role of interferon gamma (IFNγ) in mediating hemophagocytic lymphohistiocytosis, a condition characterized by accelerated erythrophagocytosis, which is also often observed in sepsis and infection, we examined IFNγ levels in the plasma after the administration of CpG-carrying RBCs ^12,14^. Plasma IFNγ and IL-6 were increased at 6 hours post-infusion of CpG-treated, but not PBS-treated, RBCs (Fig. 4I and J). Collectively these data demonstrate that CpG-carrying RBCs undergo accelerated erythrophagocytosis and initiate local and systemic immune responses.

**Table 2:** Sepsis population stratified by clinical anemia threshold (Hb ≤ 7). Data are shown as number observed (%) for categorical data, or by mean ± standard deviation or median (interquartile range) for continuous data, as appropriate to the data distribution. Comparisons shown between anemic and non-anemic subjects are by chi square tests (categorical data), Student’s T test (continuous, normally distributed), or Ranksum testing (continuous, non-normally distributed).

|  | **Anemic $Hb \leq 7$**<br>**(n=9)** | **Not Anemic $Hb > 7$**<br>**(n=39)** | **p-value** |
| --- | --- | --- | --- |
| Age | 59.3 ± 16 | 56.9 ± 16 | 0.82 |
| Female sex | 2 (22%) | 13 (33%) | 0.52 |
| Race |  |  |  |
| African American | 2 (22%) | 14 (36%) | 0.26 |
| Caucasian | 6 (67%) | 23 (59%) |  |
| Asian | 0 | 1 (3%) |  |
| Other or More than 1 | 1 (11%) | 1 (3%) |  |
| Total RBC-bound DNA ($\times 10^7$) | 6.52 (4.75 – 11.6) | 1.42 (0.70 – 3.67) | 0.0053 |
| RBC-bound 16S DNA ($\times 10^7$) | 1.21 (0.27 – 3.47) | 0.24 (0.03 – 0.80) | 0.036 |
| RBC-bound- mitochondrial DNA ($\times 10^7$) | 4.06 (0.98 – 6.32) | 0.90 (0.63 – 1.97) | 0.043 |

### Erythrophagocytosis of CpG-RBCs and CpG-induced inflammation are dependent on RBC-TLR9

The antibody mIAP301 blocks the antiphagocytic CD47 epitope on murine erythrocytes ^35^. We thus asked whether CpG-treatment would lead to a loss of CD47 detection using this antibody. CpG-treatment of erythrocytes from wild type (WT), but not TLR9 knock-out (KO) mice, resulted in a TLR9-dependent loss of CD47 detection (Fig. 5A and B). Interestingly, TLR9 KO mice also exhibited a higher number of CD47 negative RBCs in the circulation. Recent studies have demonstrated a role for monocyte/macrophage TLR9 signaling in erythrophagocytosis, these findings may reflect decreased clearance of CD47 negative RBCs in TLR9 KO mice; alternatively, these findings may reflect fundamental differences in TLR9 KO and WT erythrocytes ^12-14^.

**Figure 5.**
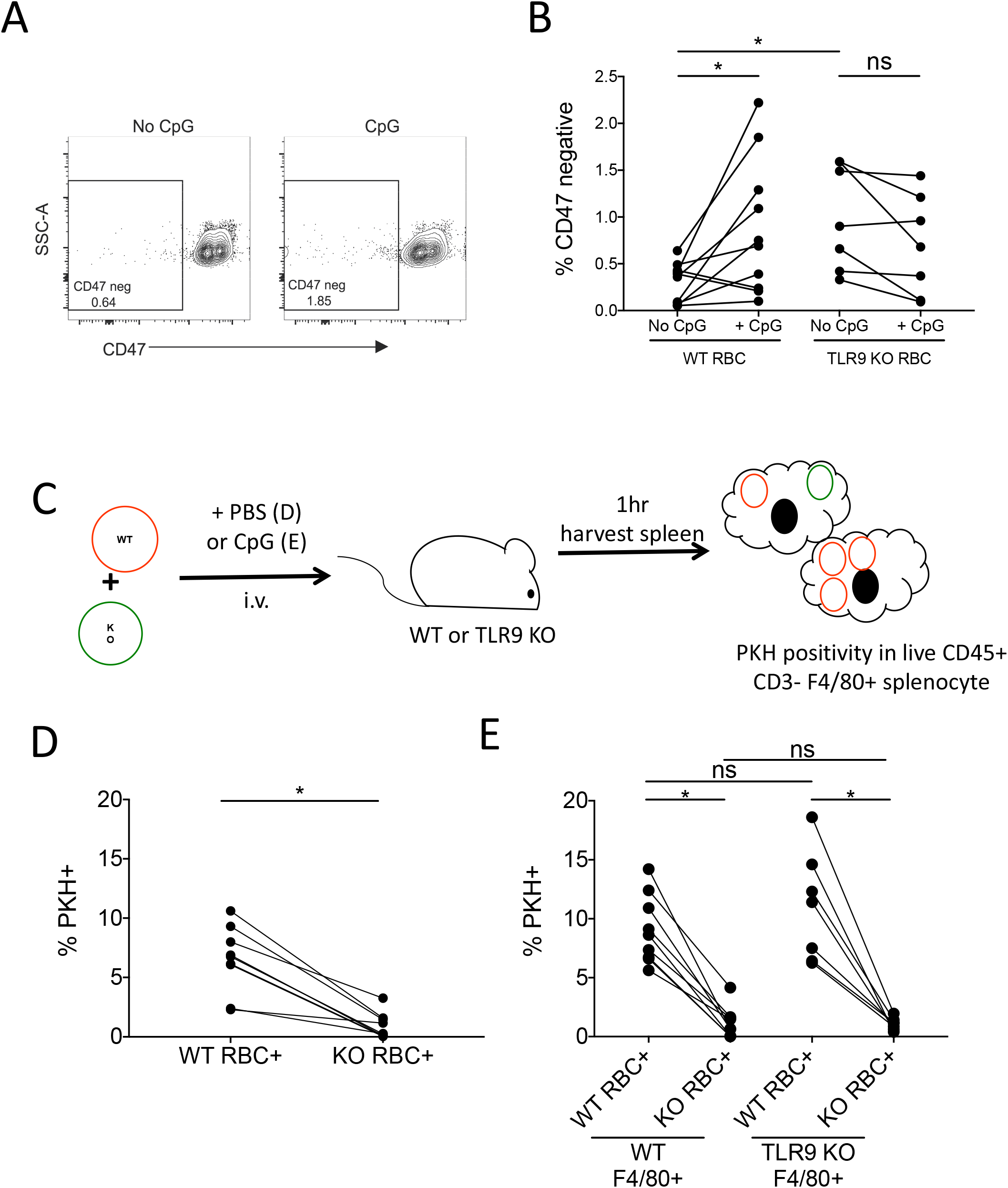
RBC-TLR9 mediates accelerated erythrophagocytosis following CpG. (**A**) CD47 detection (clone mIAP301) on C57/B6 murine RBC treated with 100nM CpG, n =10, one representative study shown. (**B**) CD47 detection on RBCs from *ex vivo* CpG-treated WT or TLR9 KO RBCs RBCs. Each line represents RBCs from an individual mouse. For WT, two-tailed paired t-test *\*P*=0.022. For KO, not significant. For non-CpG treated WT v TLR9 KO, *\*P*=0.013 Mann-Whitney U test (n=6-9 mice/group, 3 independent experiments). (**C**-**E**) WT and TLR9 KO mice were separately labeled with PKH dyes and 1×10^8^ cells from each were mixed and treated with PBS or 25nM CpG immediately before infused into WT or TLR9 KO mice. Erythrophagocytosis was assessed as PKH positivity in CD45+ CD3-F4/80+ cells. Experimental schematic is shown in (C). (D) PBS-treated RBCs in WT mice and (E) CpG-treated RBCs in WT or KO mice, n > 6, \*\*\*\**P*<0.0001, ns - not significant. Deletion of macrophage TLR9 does not alter erythrophagocytosis, results from 3 independent experiments.

To determine if accelerated erythrophagocytosis was dependent on RBC-TLR9, we examined levels of erythrophagocytosis of untreated WT and TLR9KO RBCs in naïve mice. WT and TLR9 KO RBCs were labeled with PKH dye and mixed before transfusion to naïve WT mice. Splenic phagocytes ingested more WT RBCs than TLR9 KO RBCs (Fig. 5C and D). This observation was consistent with our previous observations of higher levels of endogenous mtDNA on WT RBCs than TLR9 KO RBCs. We next asked whether TLR9 KO RBCs underwent accelerated erythrophagocytosis following CpG treatment. TLR9 KO and WT RBCs were labeled with PKH dye and combined prior to incubation with CpG. The CpG-treated RBCs were administered to WT or global TLR9 KO mice. As seen in figure 5E, F4/80+ macrophages ingested significantly higher amounts of WT than KO RBCs. This effect appears to be driven by the RBCs as macrophages from TLR9 KO mice ingested similar numbers of RBCs as WT macrophages.

To understand the role of erythrocyte TLR9 in the innate immune response *in vivo*, we generated erythrocyte TLR9 KO mice (Fig. 6A-B and table 3), and erythroid cells from erythrocyte^*tlr9-/-*^ mice (Ery^*tlr9-/-*^) did not express TLR9. To better define the role of cell-free nucleic acid-sensing and RBC-TLR9 in innate immunity, we subjected WT and Ery^*tlr9-/-*^ mice to a reductionist model of CpG-induced inflammation. CpG administration led to decreased circulating white blood cells and increased spleen weight in both WT and Ery^*tlr9-/-*^ mice (fig. S7B and C). IL-6 production in the spleen, liver, and plasma was attenuated in Ery^*tlr9*-/-^ mice (Fig. 6C-E). Ery^t*lr9*-/-^ mice also displayed attenuated liver TLR9 upregulation, endothelial activation, and IL-10 production (fig. S7D-E). Collectively, these findings suggest that RBC-TLR9 mediated CpG delivery regulates local and systemic IL-6 production.

**Table 3.** Baseline hematology labs from WT and Erythroid^*tlr9-/*^-mice. All data are presented as mean (standard deviation). Significance was determined using two tailed t-test for all parameters.

|  | **C57Bl/6J**<br>**(n=8)** | **Erythrocyte<sup>thr9-/-</sup>**<br>**(n=5)** | **p- value** |
| --- | --- | --- | --- |
| WBC (10/uL) | 458.0 (165.9) | 552.6 (234.9) | NS |
| RBC (10<sup>4</sup>/uL) | 875.3 (83.9) | 842.8 (126.9) | NS |
| HGB (g/L) | 134.3 (10.5) | 135.8 (16.4) | NS |
| HCT (10<sup>-1</sup>%) | 442.1 (41.8) | 432.6 (70.5) | NS |
| MCV (10<sup>-1</sup>fL) | 505.5 (13.6) | 512.4 (10.9) | NS |
| MCH (10<sup>-1</sup>pg) | 153.8 (4.9) | 161.8 (6.3) | p=0.026 |
| MCHC (g/L) | 304.1 (5.8) | 316.0 (16.3) | p=0.081 |
| PLT (10<sup>3</sup>/uL) | 987.8 (259.7) | 1214.2 (301.5) | NS |
| RDW-SD (10<sup>-1</sup>fL) | 238.4 (16.7) | 248.2 (10.2) | NS |
| RDW-CV (10<sup>-1</sup>%) | 137.0 (8.2) | 141.2 (8.3) | NS |
| PCT (10<sup>-2</sup>%) | 72.4 (17.6) | 89.6 (22.6) | NS |

**Figure 6.**
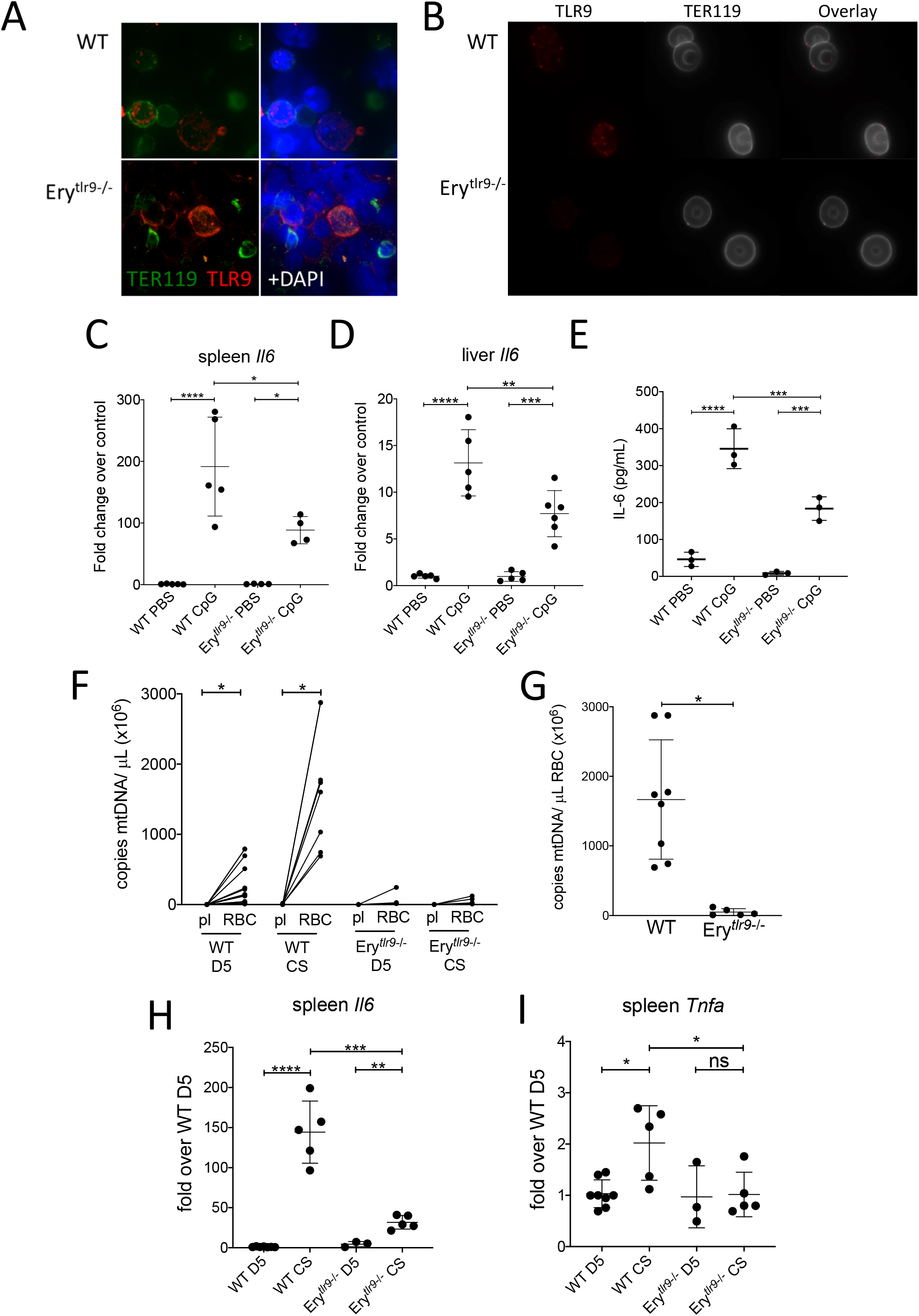
Deletion of Erythrocyte TLR9 alters host immune response. (A-B) TLR9 staining in (A) bone marrow smear and (B) mature RBC of WT and Ery^tlr9-/-^ mice. The erythroid marker TER119 is used to identify erythroid cells. Representative image from 3 individual mice were shown. (**C-E**) Inflammatory response of Ery^*tlr9-/-*^ mice following t.v. CpG for 6hr, n=3-5, one-way ANOVA with Tukey’s multiple comparison \**P*<0.05, \*\**P*<0.005, \*\*\**P*<0.0005, \*\*\*\**P*<0.0001. Quantification of *Il6* transcripts in (C) spleen and (D) liver. (E) plasma IL-6 level. (**F**-**I**) Inflammatory response of Ery^*tlr9-/-*^ mice upon cecal slurry-induced sepsis (F) mtDNA in RBC and plasma (pl) of D5- and CS-treated mice \**P*<0.001 (RBCs v plasma) for both control and CS treated mice. (G) mtDNA on RBC in cecal slurry-injected mice, \**P*=0.002. Quantification of (H) *Il6* and (I) *Tnfa* transcripts in spleens of injected mice, n=3-8, one-way ANOVA with Tukey’s multiple comparison. \**P*<0.05, \*\**P*<0.005, \*\*\**P*<0.0005, \*\*\*\**P*<0.0001, ns= not significant.

To determine whether RBC-nucleic acid-binding contributes to innate immune responses during sepsis, we subjected WT and Ery^*tlr9*-/-^ mice to a cecal slurry model of sepsis. Cecal slurry injection led to increased RBC-mtDNA sequestration in WT mice but not Ery^*tlr9-/-*^ mice (Fig. 6F and G). Because our data demonstrate accelerated clearance of DNA-bound RBCs and increased erythrophagocytosis in the spleen, we examined inflammatory responses in the spleen. Spleen IL-6 and TNFα production was attenuated in the absence of RBC TLR9 during cecal-slurry induced sepsis (Fig. 6H and I). These findings suggest that RBCs acquired CpG-DNA during sepsis, and RBC-TLR9 dependent DNA delivery drives local innate immune responses during sepsis.

## Discussion

In this study, we identify a new role of RBCs in the immune response to infection. We show that RBCs express surface TLR9 that can bind CpG-containing cell-free DNA and that both RBC TLR9 expression and RBC bound CpG are elevated in human sepsis. Under basal conditions with low cell-free DNA levels, RBCs bind CpG DNA and act as a “sink” without undergoing overt morphological changes ^4^. However, when there are high levels of plasma CpG DNA, such as during sepsis, pneumonia, or malarial infection, TLR9-dependent CpG DNA binding leads to fundamental alterations of RBC morphology, functional loss of CD47 on a subset of RBCs, accelerated erythrophagocytosis, and innate immune activation driven by RBC-DNA delivery with clearance.

Our data show that DNA binding to RBCs results in accelerated erythrophagocytosis. Surprisingly, CpG-binding to RBCs was sufficient to cause accelerated clearance in naïve mice. This may have implications for malaria pathogenesis since excess parasite DNA in the plasma of infected individuals may result in the removal and destruction of both infected and uninfected RBCs. Indeed, a recent study of *Plasmodium berghei*-infected mice showed that disruption of CD47-Sirpα binding led to accelerated erythrophagocytosis, thus implicating this interaction in the host response to malaria ^36,37^. Similarly, *Plasmodium* DNA and TLR9 have been shown to promote autoimmune anemia through a T-bet+ B cell-mediated anti-erythrocyte antibody production in *Plasmodium yoelli*-infected mice ^38^. Malarial anemia and non-malarial infectious anemia may also arise by developing inflammatory hemophagocytes or the generation of other erythrophagocytic macrophage subsets. In mice, CpG-TLR9 interactions have been shown to promote inflammatory anemia during *P. yoelli* blood stage infection, and nucleic acid-sensing TLRs promote anemia in a hemophagocytic lymphohistiocytosis model ^13,14^. Anemia is common in sepsis, and accelerated clearance of erythrocytes is a defining feature of anemia during poorly understood sepsis. Our findings provide one mechanism by which RBCs undergo accelerated clearance in sepsis as we observed RBC-TLR9 dependent accelerated clearance of CpG-carrying RBCs *in vivo*.

Accordingly, targeting RBC-TLR9 with blocking antibodies or antagonistic small molecule inhibitors may be a viable option to combat inflammatory anemia. This could potentially eliminate the enhanced CpG-TLR9 mediated RBC phagocytosis without interfering with CpG-TLR9 signaling in classical immune cells essential for host defense.

In addition to ingestion by macrophages, CpG-carrying RBCs may also be taken up by dendritic cells or other antigen-presenting cells, which may alter antigen presentation and acquired immune responses. Previous studies demonstrated that even a small fraction (0.5%) of CD47-negative RBCs could activate splenic dendritic cells and CD4+ T cells ^39-41^. Our findings of increased interferon production in mice receiving CpG-RBCs suggests that RBCs can present CpG to immune cells. Indeed, we detected CpG-containing mtDNA on RBCs during parasitic infection, pneumonia and polymicrobial sepsis. Furthermore, CpG-carrying RBCs induced an innate immune response in naïve mice as the administration of CpG-carrying RBCs led to transcriptomic changes in the spleen characterized by upregulation of host response to virus, innate immune response, and interferon signaling pathways.

In the absence of RBC-TLR9, CpG-induced IL-6 was attenuated in local and systemic compartments in a model of CpG-induced inflammation. However, in a murine model of polymicrobial sepsis, RBC-TLR9 regulated only spleen IL-6 production. Thus, how RBC-TLR9 contributes to immune dysregulation may be context-dependent, depending on the inciting stimuli. RBC-TLR9 may serve as a delivery mechanism of CpG in sterile inflammation to driving local and systemic IL-6 production. Although beyond this study’s scope, analysis of the erythrophagocytic cells in the spleen, liver and bone marrow on a single-cell level will be critical in elucidating the exact mechanisms of innate immune regulation by CpG-carrying RBCs.

TLR9 is expressed on nucleated erythrocytes in other vertebrates, including fish ^42^. Birds express the avian homolog of TLR9, TLR21, on their erythrocytes ^42^. Here, we demonstrate the presence of TLR9 and DNA binding by human, chimpanzee and murine RBCs and show a role for RBCs in sensing CpG DNA, a potent activator of the innate immune system. Humans produce over 2 million RBCs each second and are at risk for exposure to large amounts of DNA during mitophagy and nuclear expulsion. Thus, it is tempting to speculate that TLR9 is retained on erythrocytes to protect RBCs during erythroid maturation by scavenging mitochondrial DNA that escapes mitophagy. Indeed, recent studies have shown that loss of mitophagy leads to RBC destruction and anemia, and other studies have demonstrated that mtDNA that escapes mitophagy leads to cell-autonomous TLR9-mediated inflammation ^43,44^. Alternatively, given the inflammatory response observed following administration of CpG treated RBCs to naïve mice it is plausible to speculate that retention of TLR9 on RBCs promoted host survival by allowing for propagation of local signals remotely and early innate detection of cell-free DNA released during infection or following trauma. Although further studies will be required to elucidate the potential role of TLR9 in erythroid development, our findings of RBC mediated nucleic acid sensing confirm a role for TLR9 on mature erythrocytes in regulating the immune response during acute inflammation.

Consistent with our discovery of RBC-immune function, nearly two decades of research has solidified the role of another enucleated cell, platelets, in innate and adaptive immunity ^18,45,46^. Future studies examining the cooperation of platelets, RBCs, and coagulation in the innate immune response will be needed to truly understand innate immunity in the vascular compartment. Considering the current COVID-19 pandemic in which disruption of vascular homeostasis and elevated plasma mtDNA is both a risk factor for severe injury and a consequence of infection, exploring RBCs’ role in immunity will be essential for an inclusive understanding of immunity in the vascular compartment^47-49^. If CpG-delivery by RBCs drives IL-6 production in various inflammatory diseases, targeting RBC-TLR9 may be an effective way to treat cytokine storm without concomitant immune suppression known to occur with monoclonal anti-cytokine antibody therapies. Alternatively, RBC-mediated CpG delivery can be exploited in the development of vaccines and immunotherapy.

Our data demonstrate that red cells serve as DNA sensors through surface expression of TLR9, which appears to be beneficial during quiescent states, where it promotes scavenging of trace levels of CpG to prevent non-specific inflammation ^4^. However, during states of excess circulating CpG, such as sepsis, binding of CpG by RBC-TLR9 leads to accelerated clearance and inflammation. While this innate immune mechanism may be beneficial in the clearance of microbial infection and damaged RBCs, CpG-binding by RBCs likely contributes to systemic inflammation and development of anemia during pathologic states where cell-free DNA is elevated. Thus, DNA recognition by TLR9 on RBCs provides bona fide evidence for red cells as immune sentinels.

## Supporting information

GO Analysis

Differentially Expressed Genes

## Acknowledgements

We thank Peggy Zhang for her excellent technical assistance. We thank Dorothy E. Loy and Dana Hodge for preparing *P. falciparum* culture. We thank Jane Fontenot, Melany Musso and Francois Villinger at the New Iberia Research Center for providing leftover blood samples from captive chimpanzees. We thank Caroline Ittner and Ariel Weissman for providing samples from sepsis patients. We thank the Shin lab (Mark Boyer, Dr. Jessica Doerner and Dr. Xin Liu for providing samples from *Legionella*-infected mice).

## Author Contributions

Experiments were conceived and designed by NSM. Experiments were performed by DK, SSM, ML SM, AV, CC, JP, and NSM. Data were analyzed by AV, AW, NSM, SSM and ML. NJM provided samples from sepsis patients, CH, SS, SRR, BHH and AOJ provided vital reagents. GSW, CH and NSM wrote the paper.

## Data and materials availability

All data is available in the main text or the supplementary materials.

## Supplementary Materials

Materials and Methods

Figs. S1 to S7

Tables S1 and S2

Data S1 and S2

