## Supplement for "Red Blood Cells Function as DNA Sensors"

**Short Title:** RBC DNA Sensing Alters RBC Survival and Immunity

**This PDF file includes:**

Materials and Methods
Figs. S1 to S7
Tables S1 and S2

**Other Supplementary Materials for this manuscript include the following:**

Data S1 Differentially expressed genes
Data S2 GO analysis

### Materials and Methods

**Study approval.** Animal studies were conducted in accordance with the Institutional Animal Care and Use Committee at the University of Pennsylvania. Studies involving human subjects were approved by the University of Pennsylvania Institutional Review Board. Healthy volunteers between ages of 18 and 65 years gave written informed consent prior to inclusion.

**Sepsis Cohort.** RBCs were obtained from patients enrolled in the Molecular Epidemiology of Severe Sepsis in the ICU cohort (MESSI) study at the University of Pennsylvania (*49*). Patients were eligible if they presented to the medical intensive care unit (MICU) with strongly suspected infection, at least 2 systemic inflammatory response syndrome (SIRS) criteria, and evidence of new end organ dysfunction in accordance with consensus definitions by the American College of Chest Physicians. Exclusion criteria included a lack of commitment to life sustaining measures, primary reason for ICU admission unrelated to sepsis, admission from a long-term acute care hospital, previous enrollment, or lack of informed consent. Human subjects or their proxies provided informed consent.

**Experimental animals.** C57BL/6 animals were purchased from the Charles River Laboratories Inc. TLR9 knockout mice were produced by S. Akira and provided by Edward Behrens (Children's Hospital of Philadelphia). Mice lacking TLR9 in erythroid compartment are generated by crossing EpoR Cre (a generous gift from Dr. Ursula Klingmüller, German Cancer Research Center) and TLR9 flx (obtained from the European Mutant Mouse Archive). Genotype was confirmed through PCR amplification with the primers listed in table S1**.** All experimental procedures were performed on 8-12-week-old mice, 20-25 g in weight. Animal studies were conducted in accordance with the Institutional Animal Care and Use Committee at the University of Pennsylvania.

**RBC isolation from human and chimpanzee donors.** Whole blood was obtained from healthy volunteers in EDTA tubes and centrifuged for 10 minutes at 3,000 g. Plasma and buffy coat were removed. Packed RBC pellets were passed through a leukoreduction filter (Acrodisc White Blood Cell Syringe Filter, Pall Medical) with PBS or were isolated by magnetic-activated cell sorting with anti-Glycophorin A microbeads (CD235, Miltenyi Biotec). Filtered cells were centrifuged for 5 minutes at 800 g and supernatant was removed.

Specimens were also obtained from residual blood drawn for clinical purposes on admission to the ICU. Per clinical protocol, these samples were collected in EDTA-anticoagulated tubes, centrifuged at 3,000 g within 30 minutes for plasma analysis, and then stored at 4°C. RBCs were isolated within 24 hours of collection by magnetic-activated cell sorting as described above.

Chimpanzee blood samples (5-10 ml) were obtained from captive individuals (*Pan troglodytes*) housed at the New Iberia Research Center (Lafayette, Louisiana) in 10 ml ACD collection tubes (BD Biosciences). These samples were obtained for veterinary purposes only and represented leftover specimens from yearly health examinations. Whole blood was centrifuged at 1,500 g for 20 minutes (maximum acceleration and low brake speed). Buffy coats containing leukocytes were removed and red blood cells were resuspended in their respective plasma or PBS.

Resuspended red blood cells were passed through a SepaCell R-500 II filter (Fenwal), an Acrodisc filter with Leukosorb media (PALL), or a high efficiency leukoreduction filter (Haemonetics) to remove remaining leukocytes. The red blood cells were stored in RPMI (Gibco).

**RBC isolation from murine whole blood.** Whole blood from mice was collected via intracardiac puncture and RBCs were isolated as previously described (*4*). Briefly, blood was centrifuged for 10 minutes at 1,500 g, the plasma and buffy coat were removed, and packed RBC pellet was passed through a leukoreduction filter (Acrodisc White Blood Cell Syringe Filter, Pall Medical) with PBS. Filtered cells were centrifuged for 5 minutes at 800 g and supernatant was removed.

**Synthetic DNA binding and Flow cytometry (Human).** For synthetic DNA binding studies, 250,000 RBCs were incubated with 100 nM or 1 µM FITC-labled CpG (ODN2006, 10, 100 pmol / 250,000 cells), followed by flow-cytometric analysis of binding. For studies involving DNA pre-treatments, one million RBCs were incubated with DNA at 37°C for two hours with gentle shaking prior to antibody labelling. DNA treatments included the following: 400 nM, 4 µM CpG DNA (40, 400 pmol / $10^6$ cells, Synthesized by IDT), or 20 µM synthetic CpG DNA representing sequences from *P. falciparum* (2,000 pmol / $10^6$ cells). Cells were labeled with the following antibodies: mouse monoclonal against TLR9 (5 µg, Clone 5G5, Abcam), CD47 (0.5 µg, CC2C6, Biolegend), or CD47 (1 µg, 2D3, eBioscience).

For calcein studies, one million RBCs were first labeled with calcein-AM (5 µM, Fischer) prior to treatment with 8 µM of CpG DNA (800 pmol / $10^6$ cells). For phosphatidylserine externalization studies, 250,000 RBCs were pre-treated with 100 nM or 1 µM CpG (10, 100 pmol / 250,000 cells) prior to labeling with Annexin V (Life Technologies) in Annexin V Binding Buffer (Life Technologies). FACS acquisition and analysis was performed using the LSR Fortessa (BD Biosciences) and FlowJo Software.

**Bacterial DNA binding to human RBCs.** *S.aureus* bacteria (25923D-5) and *P. aeruginosa* genomic DNA (47085DQ) were obtained from ATCC. Bacterial culture were gifts from Dr. Sunny Shin (*L. pneumophila*, University of Pennsylvania), Dr. Hao Shen (*S. pneumoniae*, University of Pennsylvania), and Dr. G Scott Worthen (*K. pneumonia,* University of Pennsylvania). Bacterial genomic DNA was isolated and purified from bacteria using DNeasy DNA blood & tissue kit (Qiagen). $1x10^7$ RBCs were incubated with 0ng (PBS), 0.1ng, 1ng, or 10ng purified DNA in 2hr at 37°C in gentle shaking in DNA LoBind tubes (Eppendorf) and washed with PBS twice. RBC-associated DNA was extracted from samples using DNeasy kit (Qiagen), and genomic DNA were quantified with qPCR using primer or probes in table 4. In addition, the presence of 16S DNA on *L. pneumophila* DNA-treated RBC were quantified with 16S primer in table S1 and the corresponding amplicons were visualized with agarose gel.

***P. falciparum* culture.** *P. falciparum* cultures were propagated as previously described and were provided by Drs. Odom John and Hahn (University of Pennsylvania) (*50*). Parasites were incubated with 4% packed red blood cells (BioIVT) in RPMI media with glutamine supplemented with 25 mM HEPES, 0.5 g/100 ml Albumax-II, 0.36 mM hypoxanthine, and 0.01 mg/ml gentamycin and cultured in an atmosphere of 90:5:5 N:O2:CO2 gas. Culture media was spun for 5 minutes at 1,500 rpm prior to incubation with RBCs.

**Malarial DNA binding by RBCs.** RBCs ($10^6$, $10^7$, $10^8$) isolated from healthy volunteers were incubated with 200 µl PBS or culture media from *P. falciparum* cultures at 37°C for two hours with gentle shaking in DNA LoBind tubes (Eppendorf). Following incubation, RBCs were added to 500 µl of 20% sucrose and centrifuged for 3 minutes at 13,000 rpm to separate the supernatant from the intact RBCs. DNA was extracted from samples using a commercially available kit (DNeasy, Qiagen). PCR amplification of a 500 bp fragment of the mitochondrial cytochrome c oxidase subunit III (*coxIII*) gene of *P. falciparum* was performed using primers listed in table S1. Amplicons were verified by gel-purification and sequencing by the University of Pennsylvania Genomics Analysis Core.

**Synthetic DNA binding and Flow Cytometry (Mouse).** RBCs were isolated as described above and $10^6$ RBCs were labeled with monoclonal TLR9 (5µg, Clone 5G5, Abcam). In separate studies RBCs ($10^6$) were incubated with 400 nM or 4 µM CpG (ODN1826) DNA (40 or 400 pmol / 1e6 cells) (IDT) at 37°C for two hours with gentle shaking. RBCs were then labeled with PE CD47 (1µg, clone miap301, Biolegend). FACS acquisition and analysis was performed using the LSR Fortessa (BD Biosciences) and FlowJo Software.

**Detection of mtDNA and 16s in human samples**. DNA was extracted from $10^7$ RBCs using a commercially available kit (DNeasy, Qiagen). mtDNA in samples was quantified by amplifying amplicon corresponding to human *CoxI* using SYBR green I (Roche) and qPCR assay (Life Technologies). 16S DNA in samples was quantified using Taqman™ Fast Universal PCR Master Mix (Applied Biosystems). Primer and probe sequences are described in table S1.

**Measurement of mtDNA on murine plasma and RBCs.**

Whole blood from mice infected with *Legionella pneumophila* or *Toxoplasma gondii* were obtained from collaborators at the University of Pennsylvania (Sunny Shin Lab and Chris Hunter Lab respectively). Methods for each individual infection are previously described (*51, 52*).

Whole blood was centrifuged for 5 minutes at 10,000 rpm. 5 µl aliquots of plasma and RBCs were obtained for DNA extraction with DNeasy blood & tissue kit (Qiagen), followed by mtDNA quantification with SYBR green I (Roche) using primers in table 4.

**Cecal Surry Model**
Donor mice were euthanized with ketamine/xylazine (80/10 mg/kg), and sterile instruments were used to open the abdominal cavity and remove the cecum. One end of the cecum was cut and cecal contents were pushed out into a pre-weighed tube. Cecal contents from all donor mice were pooled, and ice-cold 5% dextrose in water (D5) was added to obtain a final concentration of 200mg/ml cecal slurry. Aliquots of cecal slurry were frozen at -80°C. For experiments, aliquots of cecal slurry were thawed under warm water and immediately used for injection. Intraperitoneal injections of 2 mg cecal content per gram of recipient mice were administered to mice. Body temperature of mice were monitored every 2hr for signs of distress and disease. Mice lacking a physiologic response to cecal slurry (i.e. no hypothermia) were excluded from subsequent studies. At 6hr post-injection, mice were sacrificed with ketamine/xylazine (80/10 mg/kg), and whole blood was obtained via intracardiac puncture. Mice were then cervically dislocated and tissues were weighted, harvested, snap frozen in liquid nitrogen, and stored at -80°C until usage. Whole blood was centrifuged for 5 minutes at 10,000 rpm, and 5µl aliquots of plasma and RBCs were obtained. DNA was extracted from RBC and plasma samples, mtDNA quantification was performed as described above.

**Electron microscopy**. RBCs ($10^5$) isolated from healthy volunteers were treated with 100 nM Class B, 2006 CpG DNA (10 pmol /$10^5$ cells) at 37°C for two hours with gentle shaking. Following incubation, cells were washed with PBS and fixed with 0.05% glutaraldehyde

(Polysciences). Scanning Electron Microscopy sample preparation and acquisition was performed by the University of Pennsylvania Electron Microscopy Resource Laboratory. Cells were examined with a JEOL 1010 electron microscope fitted with a Hamamatsu digital camera and AMT Advantage image capture software.

**Confocal microscopy.** RBCs ($10^5$) isolated from healthy volunteers were incubated with 100 nM Class B, 2006 CpG DNA (10 pmol /$10^5$ cells) or 2336 GpC control DNA (10 pmol/$10^5$ cells) at 37°C for two hours with gentle shaking. For F-actin, Spectrin, and Band3 staining, RBCs were then fixed using 0.05% glutaraldehyde, permeabilized with 0.1% Triton-X, washed, and stained (F-actin: 5 µg/ml, ab130935, Abcam; Spectrin: 1:100 dilution, ab2808, Abcam; Band3: 10µg/ml, sc133190, Santa Cruz). For TLR9 surface staining, RBCs were first labeled with the membrane dye PKH (MIDI26-1KT, Sigma) and then washed and fixed with 0.05% EM grade glutaraldehyde prior to the addition of TLR9 antibody (20µg/ml, Clone 5G5, Abcam). For intracellular staining of chimpanzee RBCs, RBCs were fixed, permeabilized, washed, and stained with mouse anti-TLR9 (5 µg/ml, 26C593.2, Novus Biologicals) in addition to PKH dye. For CD47-TLR9 colocalization staining, RBCs were fixed, permeabilized, washed, and stained with mouse CD47-2D3 (10 µg/ml, EBioscience) and rabbit TLR9 (10 µg/ml, ab187148, Abcam). Confocal images were acquired using the SCTR Leica Confocal microscope, and images were analyzed using ImageJ software.

**Imaging flow cytometry**. For TLR9 staining and CpG binding, RBCs ($5x10^5$) isolated from healthy volunteers were incubated with varying doses – 1 µM (100 pmol / $5x10^5$ cells) , 500 nM (50 pmol / $5x10^5$ cells), 100 nM (10 pmol / $5x10^5$ cells), 50 nM (5 pmol / $5x10^5$ cells), 25 nM

(2.5 pmol / $5x10^5$ cells), 10 nM (1 pmol / $5x10^5$cells), or 5 nM (0.5 pmol / $5x10^5$ cells) - of Class B, 2006 CpG DNA AlexaFluor-594 (IDT) at 37°C for two hours with gentle shaking. RBCs were then labeled with FITC-conjugated mouse monoclonal TLR9 (1 µg, clone 5G5, Abcam). For TLR9-CD47 double staining and CpG binding, RBCS ($10^6$) were incubated with 400 nM Class B, 2006 CpG DNA AlexaFluor-674 (40 pmol/$10^6$ cells) (synthesized by IDT) at 37°C for two hours. RBCs were then labeled with FITC-conjugated mouse monoclonal TLR9 (1 µg, clone 5G5, Abcam) and Pacific Blue-conjugated mouse monoclonal CD47 (1 µg, clone CC2C6, Biolegend). TLR9 and CD47 expression and CpG binding to RBCs were visualized and quantified by Image Stream analysis. Raw Image Files (RIF) were analyzed in IDEAS software. The automated feature finder generated Fisher Discriminant values to differentiate sub populations of RBCs. Mean pixel and Intensity features clearly discriminated two populations of cells that exhibited different morphologies by phase contrast. The intensity feature is a measure of the fluorescence intensity of the whole RBC, while the mean pixel feature indicates the mean pixels per RBC. These subpopulations were denoted as either ‘smooth’ or ‘altered’. Imaging flow cytometry was performed by the Penn Flow Cytometry and Cell Sorting Core.

**Osmotic fragility assay and hemolysis assays.** $5x10^5$ freshly isolated RBCs from human donors were incubated with indicated concentrations of CpG in PBS for two hours at 37°C with gentle shaking at 90 rpm. Cells were pelleted and washed in 1 ml PBS and resuspended in 100 µl PBS, followed by incubation in 15 ml NaCl solutions (0.9%, 0.6%, 0.3%, 0.1%, or 0% NaCl) in deionized water at room temperature for 10 minutes. The reactions were centrifuged at 800g for 10min at 4°C on slow brake. Hemoglobin content in supernatant was determined using QuantiChrom™ Hemoglobin Assay according to manufacturer’s protocol. Hemoglobin content in the reaction of RBCs without CpG treatment and incubated with water (0.00 Osml/L) was set at 100% and data were normalized against it.

**Erythrophagocytosis studies – PKH-labeled RBC.** RBCs were obtained from WT or TLR9 KO mice through leukoreduction as described above and labeled separately with PKH26 or PKH67 dyes (Sigma). To label RBCs, $2x10^9$ RBCs were then stained with 4 µl PKH dyes per reaction for 5 min at room temperature as described in manufacturer's protocol; staining was stopped by adding excess DMEM +10% FBS for 1min. Cells were washed twice with DMEM + 5% FBS and resuspended in PBS. $1x10^8$ RBCs from each of WT or TLR9 KO mice were mixed in a final $2x10^8$ RBC/ 200 µl solution in DNA lo-bind tubes. PBS or 25 nM CpG (ODN1826) were added to RBC mixture immediately before transfusion and then transfused to WT or TLR9 KO mice through tail vein injection. At 1hr post-transfusion, mice were sacrificed with ketamine/xylazine. Spleens were harvested, weighed, minced, and passed through a 70µm nylon mesh to obtain a single-cell suspension, followed by RBC lysis by ACK buffer (Thermo Fisher Scientific) to obtain white blood cells (WBC). One million WBC were stained for 30min at 4C for each of the followings: Live/Dead Fixable viability violet stain, Fc block (CD16/32, clone 93), and antibody cocktail against F4/80 (AlexaFluor700, BM8), CD45 (APC, 30-F11), and CD3 (BV711, 145-2C11). At least 10,000 live cells events were analyzed with an LSR Fortessa (BD Biosciences). Gating analysis was performed using FlowJo software (FlowJo, LLC, Ashland, Oregon).

**Erythrophagocytosis studies – GFP-expressing RBC.** GFP-positive RBCs obtained from GFP-expressing mice (C57BL/6-Tg(UBC-GFP)30Scha/J from Jackson Laboratories) were incubated with CpG (50 µg / 100 µl packed RBCs) at 37°C for two hours with gentle shaking in DNA- Lo Bind tubes (Eppendorf). Following incubation and subsequent washes, 200 µl GFP RBCs and DNA-treated GFP RBCs were transfused to C57BL/6 mice through retro orbital injection (45 minutes before transfusion, 200 µl of blood was removed from each mouse). One hour following injection, mice were sacrificed with isoflurane and cervical dislocation. Spleens were harvested and processed as above. For flow staining, cells ($10^6$ cells) were resuspended in 100 µl staining buffer (PBS+0.1% sodium azide) and incubated with anti-mouse CD16/32 antibody (Fc block, eBiosciences, San Diego, CA) for 10 min at 4°C to block nonspecific binding. Cells were then stained with the following antibodies (F4/80 PE, #123109, Biolegend; CD11b eFluo450, M1/70, #101211, Biolegend; CD45 APC, clone 30-F11, #559864, BD Pharmingen; CD3 BUV395, 145-2C11, #564661, BD) or appropriate isotype controls (0.25–1.5 µg/$10^6$ cells) for 30 minutes at 4ºC. Cells were then spun and resuspended in staining buffer for viability staining (Fixable Viability Dye, eFluor780, #65-0865-14, eBioscience) for 30 minutes at 4ºC. Cells were fixed in 2% paraformaldehyde for 10 minutes, spun and resuspended in 500 µl PBS. At least 50,000 CD45+ cells were analyzed with an LSR Fortessa (BD Biosciences). Gating analysis was performed using FlowJo software (FlowJo, LLC, Ashland, Oregon).

**Bone marrow smear and staining.** Mice were sacrificed with intraperitoneal injections of ketamine/xylazine (80/10 mg/kg), followed by cervical dislocation. Using sterile scissors, tibia bones from were separated from muscles and cut at the proximal end. A PBS-moistened No.0 art brush was used to obtain bone marrow biopsy, which was gently and thinly brushed onto Superfrost Plus microscope slide (Fisher) and air dried prior to staining. The brush was washed by dipping in two charges of 95% ethanol, followed by three charges of PBS prior to re-usage. Air-dried smears were permeabilized in 100% methanol at -20°C for 10min, washed in wash buffer (PBS + 0.05% tween20), and blocked in blocking buffer (wash buffer + 1% BSA + 10% normal goat serum) for 1hr at room temperature and probed with antibodies against TLR9 (Abcam, ab37154) and TER119 (Biolegend, clone TER-119) or appropriate isotypes in blocking buffer overnight at 4ºC. Slides were then washed and stained with secondary antibodies derived from goat and Hoechst 33342 for 1hr at room temperature and then mounted on Fluoromount G. Images were taken on Nikon 2A microscope.

**Immunofluorescence for murine erythrocyte.** Circulatory mature RBCs were isolated as above. $2.5x10^5$ cells were used per staining reaction. RBCs were fixed in 0.05% glutaraldehyde and permeabilized with 0.1% Triton-X, washed in FACS buffer (PBS + 2% FBS), blocked in PBS + 1% BSA + 5% goat serum, and probed for TLR9 (Abcam, ab37154, 5µg/mL) and TER119 (Biolegend, clone TER-119, 5µg/mL) or appropriate controls overnight at 4ºC. Cells were then washed and stained with secondary antibodies derived from goat for 1hr at room temperature and then mounted on Fluoromount G.

**Murine CpG-RBC transfusion model.** RBCs ($10^9$) were incubated with 150 µM 1826 CpG DNA (15,000 pmol / $10^9$ cells, synthesized by IDT) at 37°C for two hours with gentle shaking. Following incubation and subsequent washes, PBS, RBCs, or CpG-treated RBCs were injected via tail vein. Six hours following injection, mice were sacrificed with intraperitoneal injections of ketamine/xylazine (80/10 mg/kg). Whole blood was obtained via cardiac puncture and centrifuged for 5 minutes at 10,000 rpm to isolate plasma. Interferon gamma and IL-6 in the plasma was measured by ELISA (R&D). Spleens were formalin fixed prior to paraffin embedding. H&E staining was performed by the Penn Veterinary School Comparative Pathology Core. Spleen H&Es were read and scored by a veterinary pathologist. Spleen injury was determined by the presence of neutrophils or red pulp congestion (0 for absent 1 for present). 3-4 mice/group from 2 independent studies is shown. RNA-sequence analysis was performed on the spleen of mice transfused with PBS-RBCs or CpG-RBCs (Genewiz). Data analysis was performed using DEseq2 (Genewiz) and EdgeR.

**Quantitative PCR for mouse tissues.** Snap frozen mouse tissues were pulverized with Bessman tissue pulverizer on dry ice. RNA was extracted from powered tissues using RNeasy Plus kit (Qiagen), followed by cDNA synthesis using SuperScriptTM First strand synthesis with random hexamer. Target genes were quantified with Taqman® assay listed in table 5. Data were normalized to 18S using ΔΔCt method and expressed as fold change ($2^{-\Delta\Delta Ct}$).

**Multiplex cytokine quantification.** Plasma cytokines were quantified using the U-plex cytokine (MSD) assay according to the manufacturer's protocol. The U-plex is a customary designed 7-plex assay for IFNγ, TNFα, MIP, IL-6, IL-10, IL-12p70, and KC.

**Immunoprecipitation of TLR9 and CD47 from human RBCs.** Protein G beads (GE) were incubated with 5 µl of mouse monoclonal TLR9 antibody (26C593.2, Abcam) or CD47 antibody (2D3, eBioscience) and incubated for three hours at room temperature. RBCs ($10^9$) were lysed in RIPA buffer (50 mM Tris, 150 mM NaCl, 1% NP40, 0.1% SDS, 0.5% sodium deoxycholate and distilled $H_2O$) for 15 minutes on ice. Lysed RBCs were added to the antibody and beads and incubated overnight at 4°C. Following centrifugation at 10,000 rpm for 20 minutes the beads were washed five times with RIPA buffer. 40 µl of sample buffer was added and then the beads were heated to 95°C for 5 minutes. Proteins were resolved by SDS-page, and immunoblotting was performed with mouse monoclonal TLR9 (Abcam), rabbit monoclonal TLR9 (Abcam), mouse BAND3 (Santa Cruz), rabbit monoclonal CD47 (Abcam), or mouse CD47 (eBioscience).

**Oligodeoxynucleotides (ODNS) utilized.**

2006 CpG (human): 5’T*C*G*T*C*G*T*T*T*T*G*T*C*G*T*T*T*T*G*T*C*G*T*T-3’

1826 CpG (mouse): 5’-T*C*C*A*T*G*A*C*G*T*T*C*C*T*G*A*C*G*T*T-3’

Malarial CpG 2 (18): 5’- T*C*G*T*C*G*T*C*G*T*C*G*T*C*G*T*C -3’

Control GpC (human): 5’G*G*G*GAGCAGCTGCTGG*G*G*G*G*G-3’

*phosphorothioate bond

**Supplemental Table 1. Primer and probe sequences for PCR and qPCR**

| Target | Forward Primer (5'->3') | Reverse Primer (5'->3') | Probe (5'->3') |
|---|---|---|---|
| *S. aureus* (16S) | TCGGMTCGTAAAACTCTGTT | CTGCTGGCACGAAGTTAGC | /56-FAM/AAGAACATA/ZEN/TGTGTAAGTAACTGTGCACA/3IABkFQ/ |
| *K. pneumoniae* (16S) | GCCTTCGGGTTGTAAAGY | CTGCTGGCACGAAGTTAGC | /56-FAM/TTAATAACC/ZEN/TYRKCGATTGACGTTACCC/3IABkFQ/ |
| *L. pneumophila* (16S) | TACCTACCCTTGACATACAGTG | CTTCCTCCGGTTTGTCAC | /56-FAM/GAGTCCCCA/ZEN/CCATCACATG/3IABkFQ/ |
| *P. aeruginosa* (16S) | CTGGAAGCAGGATGGCTATT | CAGTAGCGGGAAGAGAATGTAG | /56-FAM/AACTGCTCT/ZEN/TCCACCGACAACGAC/3IABkFQ/ |
| *S. pneumoniae* (CpsA) | GCTGTTTTAGCAGATAGTGAGATCGA | TCCCAGTCGGTGCTGTCA | /56FAM/AATGTTACGCAACTGACGAG/3IABkFQ/ |
| 16S (universal) | BSF8:AGAGTTGATCCTGGCTCAG | BSR357:CTGCTGCCTYCCGTA | /56-FAM/TAA+CA+CATG+CA+AGT+CGA/3BHQ_1/ |
| *P. falciparum* (*coxIII*) | AGCGGTTAACCTTTCTTTTTCCTTACG | AGTGCATCATGTATGACAGCATGTTTACA | N/A (gel amplicon) |
| human COXI (MT-CO1) | TGATCTGCTGCAGTGCTCTGA | TCAGGCCACCTACGGTGAA | N/A (SYBR green) |
| murine COXI (mt-Co1) | GCCCCAGATATAGCATTCCC | GTTCATCCTGTTCCTGCTCC | N/A (SYBR green) |
| EpoR Cre | GTGTGGCTGCCCCTTCTGCCA | CAGGAATTCAAGCTCAACCTCA | N/A (genotyping) |
| TLR9 flx | CGGTTAATGGTAGCACTTGG | GCTTTTGCTCAGAACACAACC | N/A (genotyping) |

**Supplemental Table 2. Taqman assay used for qPCR of murine genes**

| Target | Taqman assay system |
|---|---|
| murine *Tnfa* | Mm00443258_m1 |
| murine *Il6* | Mm00446190_m1 |
| 18S | Hs99999901_s1 |
| murine *Sele* | Mm00441278_m1 |
| murine *Tlr9* | Mm00446193_m1 |

**Statistics.** Differences between groups were compared using a t-test, Mann-Whitney U test, or one way ANOVA, as appropriate. All statistical analyses were performed using Sigma Plot 13 software (Systat Software Inc). A $p<0.05$ was considered significant for all analyses.

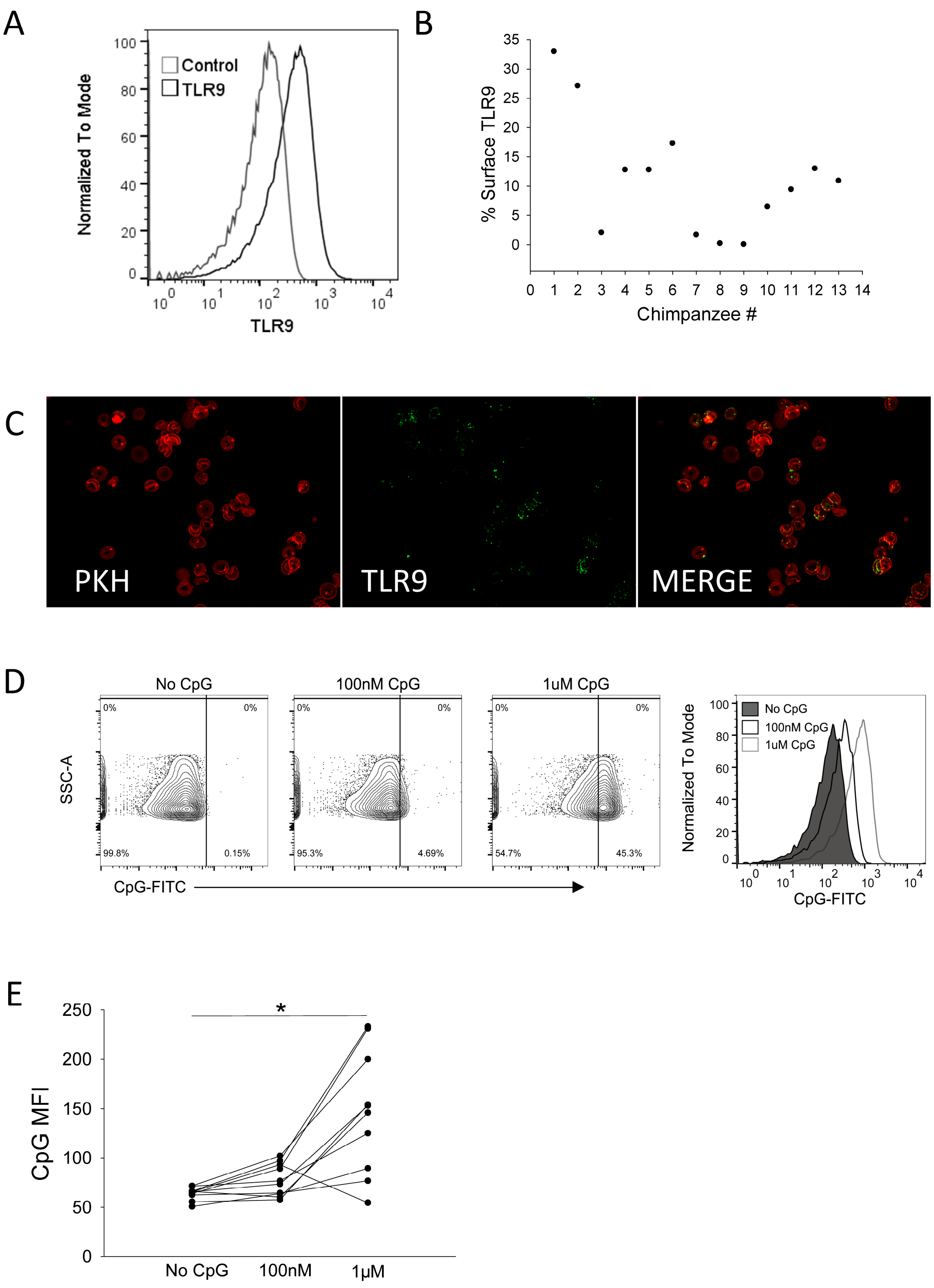


**Fig. S1. RBCs from mice and chimpanzees and mice express TLR9 on their surface.** (**A**) MFI of surface TLR9 expression on unfixed, non-permeabilized RBCs from a chimpanzee, (n=13, one representative sample shown). (**B**) Surface TLR9 expression on RBCs from each chimpanzee (n=13). (**C**). Imaging of TLR9 expression on fixed and permeabilized chimpanzee RBCs. (**D-E**) Concentration-dependent binding of CpG to Chimpanzee RBCs. (D) Representative flow plots from one chimpanzee and (E) corresponding MFI data, n=10 **P*=0.002, one-way ANOVA.

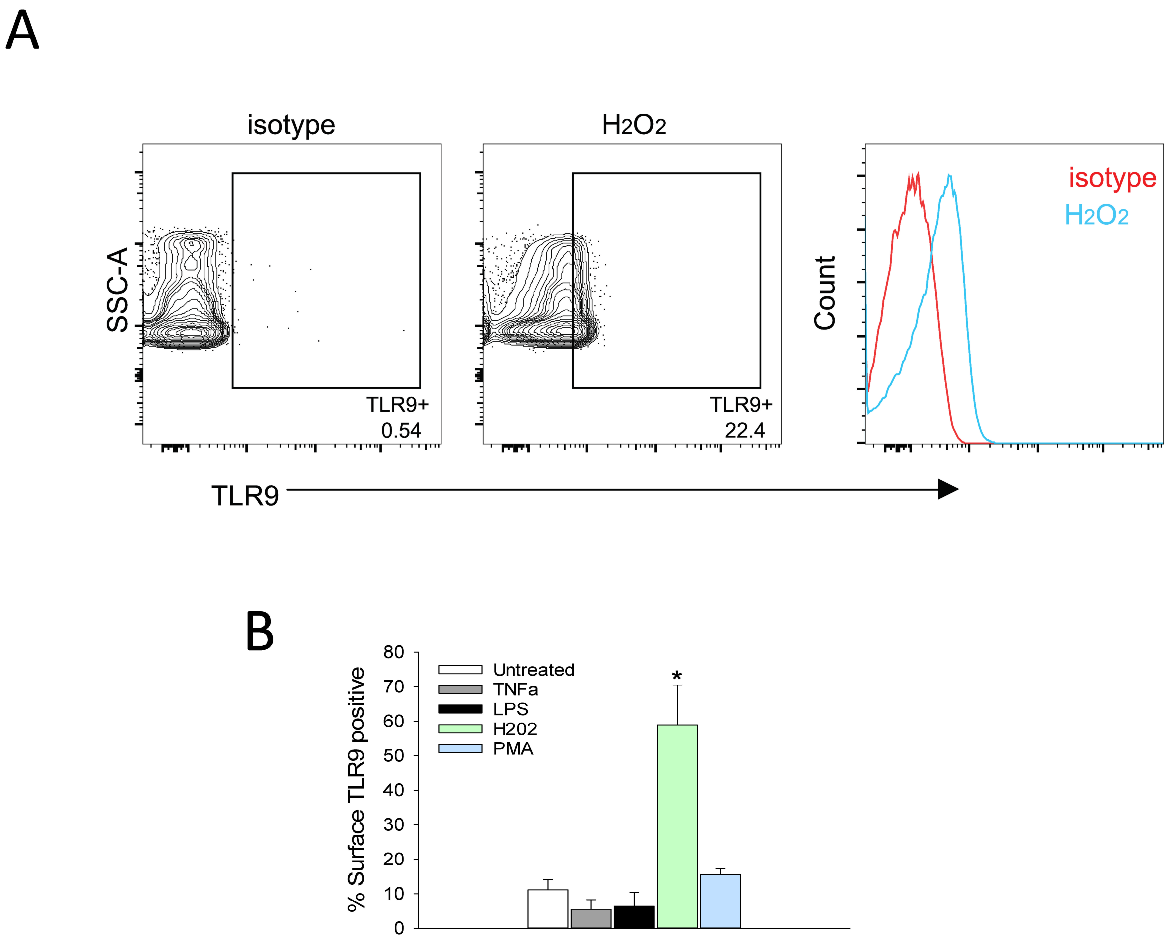


**Fig. S2. Effects of immune-activating reagents on surface TLR9 detection.** Human RBCs treated with indicated reagents were stained for surface TLR9 expression analyzed by flow cytometry, representative flow plot shown in (**A**) and quantified data in (**B**). *$P$=0.036 one-way ANOVA, Dunn's post-hoc analysis, untreated v $H_2O_2$ treatment, n=3 independent studies and RBC donors.

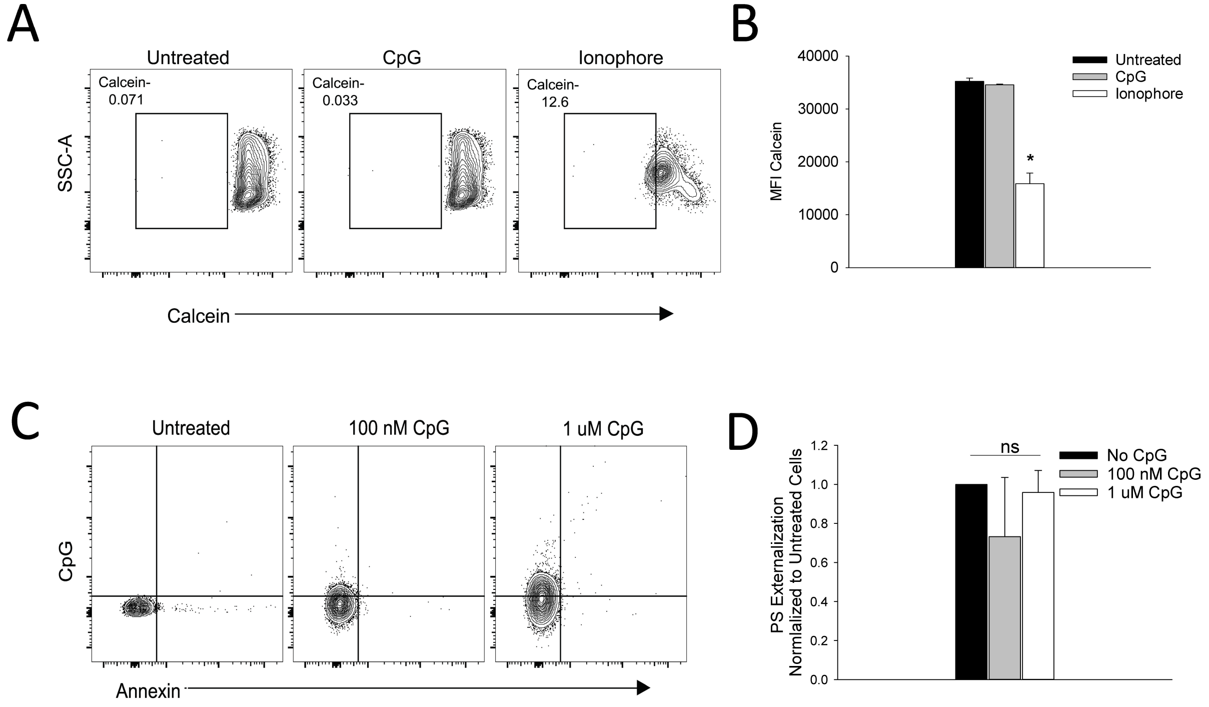


**Fig. S3. Viability and PS externalization is not altered following CpG treatment of RBCs.** (**A**) Calcein AM detection in CpG or ionophore-(A23187) treated RBCs. (**B**) Mean fluorescent intensity (MFI) of calcein AM in RBCs treated with CpG or calcium ionophore, n=3 donors, **P*<0.001, t test untreated RBC v ionophore, *P*=NS, untreated RBCs v CpG. (**C**) Flow cytometry analysis of phosphatidylserine (PS) externalization on RBCs incubated without or with fluorescent CpG (100 nM, 1 μM) prior to Annexin V staining. (**D**) Average PS externalization, normalized to untreated cells, following CpG treatment, n=4 healthy donors.

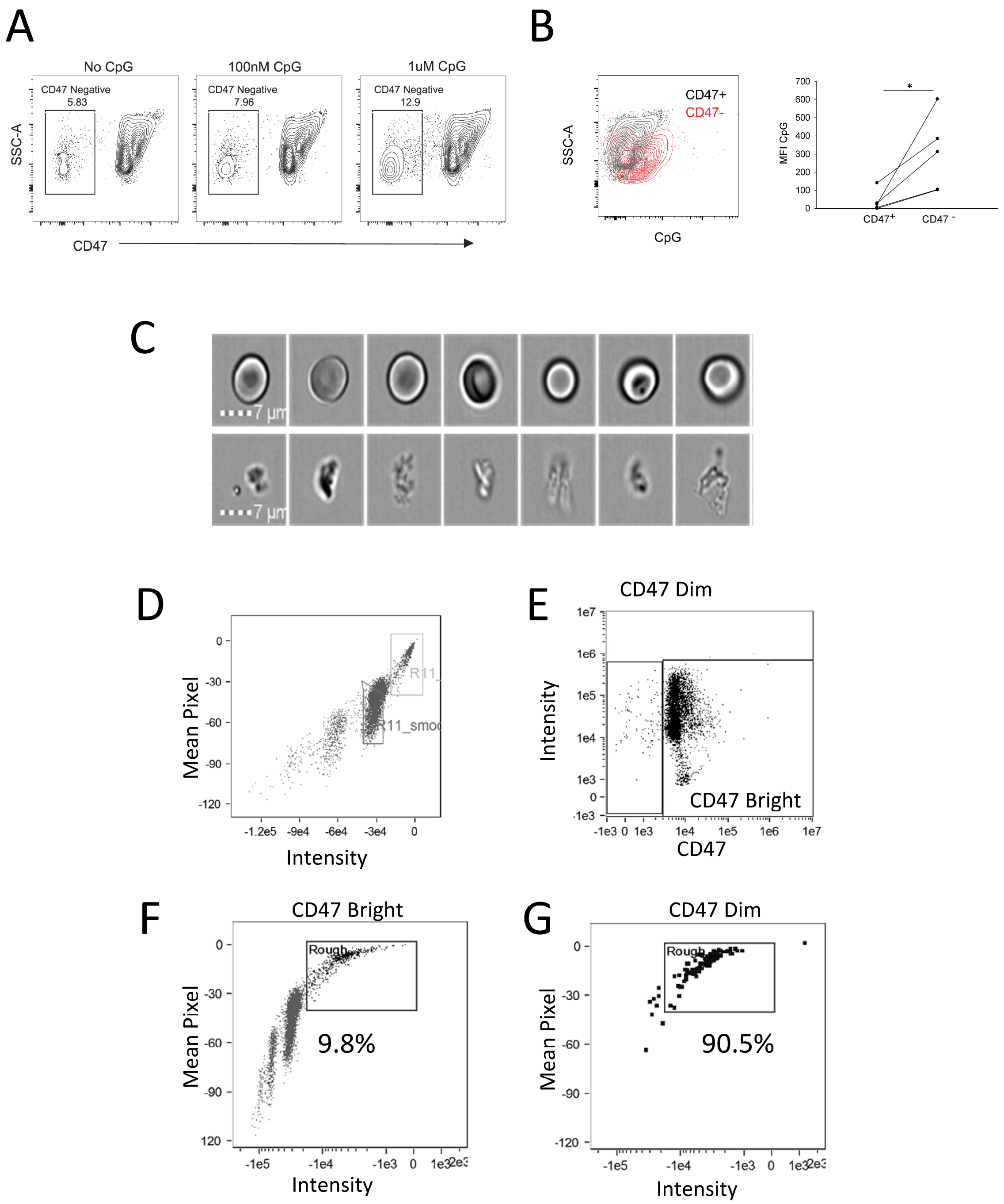


**Fig. S4. Characterization of CD47 negative and positive cells following CpG treatment.** Human RBC were incubated with CpG and analyzed by (**A**-**B**) flow cytometry and (**C**-**F**) imaging flow cytometry. (A) CpG-positivity in CD47+ and CD47- populations and the corresponding MFI data in (B). (C) Imaging flow cytometry to determine characteristics of cells following DNA incubation reveals 2 distinct populations of smooth (top panel) and rough (bottom panel) cells. (D) Gating strategy- Mean Pixel v Intensity identifies "rough" and "smooth" populations. (E) Gating for CD47 negative and positive populations. (F) The "rough" gate was applied to both CD47 Bright (left) and CD47 Dim cells (right). CD47 high cells contain 9.8% rough cells while CD47 negative cells contain 90.5% rough cells.

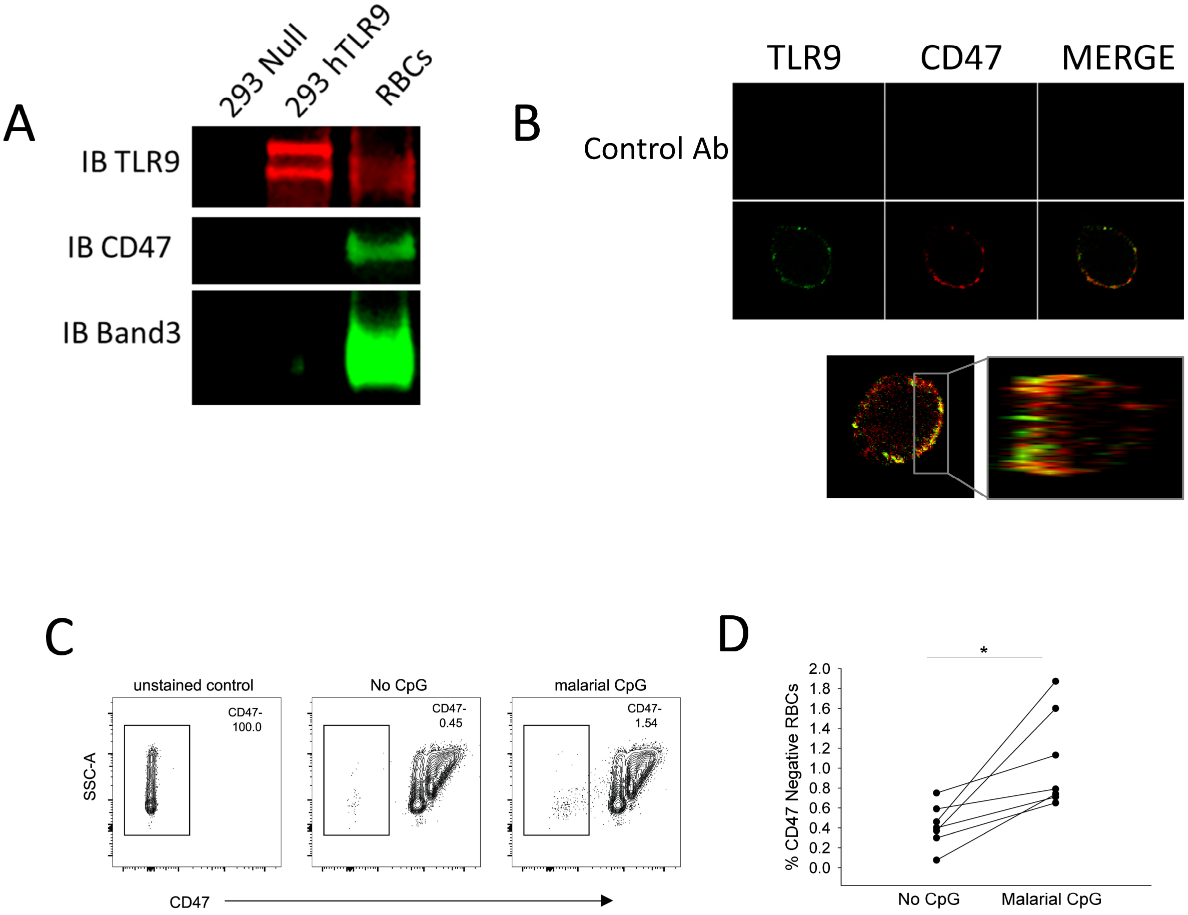


**Fig. S5. TLR9 and CD47 interact on the RBC surface.** (**A**) Immunoprecipitation of TLR9 from RBCs reveals association with Band3 and CD47, IP, immunoprecipitation; IB, immunoblot. (**B**) Confocal imaging of CD47 and TLR9, Green, CD47 (2D3), Red, TLR9 (merged and Z stack images are shown in the bottom panel). (**C**-**D**) Flow cytometry analysis of synthetic malarial CpG-treated RBCs labeled with CD47 antibodies (clone CC2C6) following CpG treatment. Gating strategy (C) and corresponding data form individual donors (D).

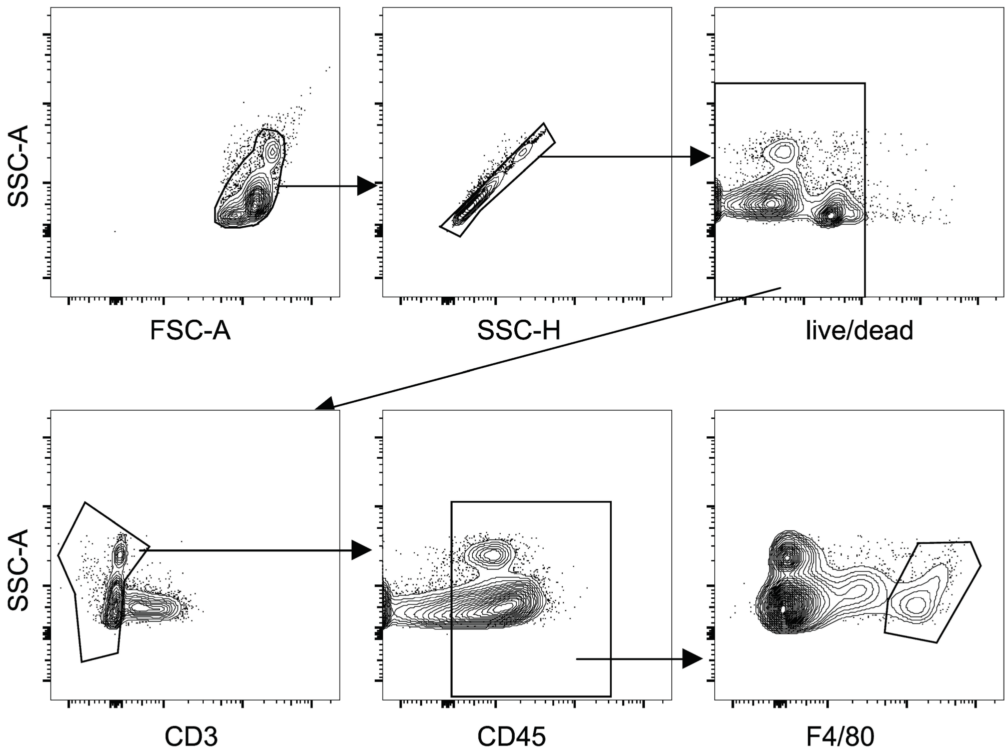


**Fig. S6. Gating strategy on mouse splenocytes to identify red pulp macrophages.** For analysis of red pulp macrophages (RPMs), splenocytes were labelled with antibodies against CD3, CD45, and F4/80. RPMs were identified live cells that were $CD3^{-}$, $CD45^{+}$, F4/80 high.

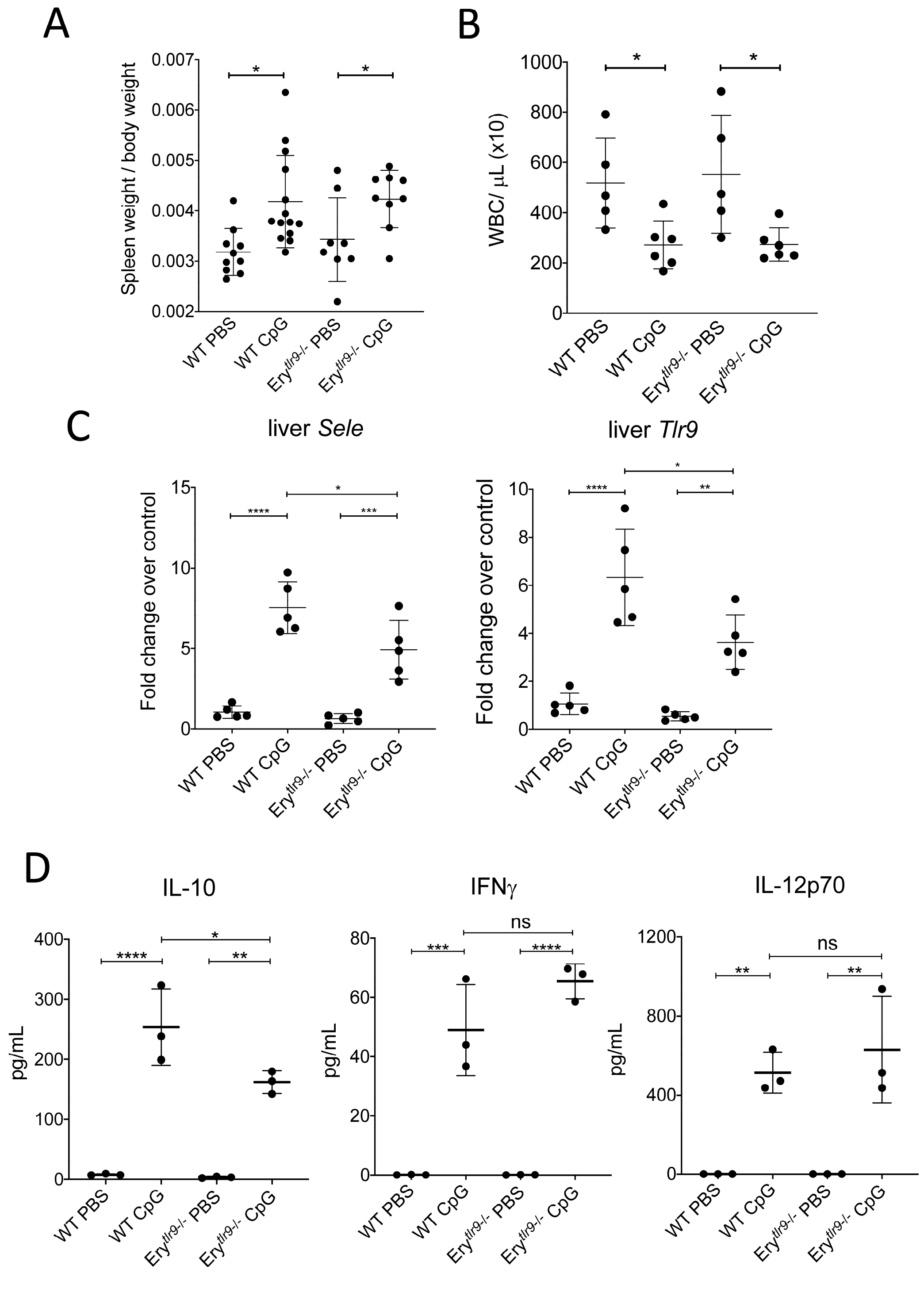


**Fig S7: Physiologic changes and cytokine response of CpG-injected erythroid TLR9-deficient mice.** WT or Ery$^{tlr9-/-}$ mice were injected with PBS or 200µL of 150µM CpG intravenously for 6hr. (A) Normalized spleen weight and (B) circulatory white blood cell counts of injected mice. (C) Measurement of *Sele* and *Tlr9* gene expression in liver by qPCR. (D) U-plex multiplex quantification of plasma cytokine levels for IL-10, IFNγ, and IL-12p70.
